# A Metabolic Complex Involved in Tomato Specialized Metabolism

**DOI:** 10.1101/2025.02.17.638719

**Authors:** Varun Dwivedi, Ernest Okertchiri, Adam Yokom, Craig A. Schenck

## Abstract

Specialized metabolites mediate diverse plant-environment interactions. Although, recent work has begun to enzymatically characterize entire plant specialized metabolic pathways, little is known about how different pathway components organize and interact within the cell. Here we use acylsugars – a class of specialized metabolites found across the Solanaceae family – as a model to explore cellular localization and metabolic complex formation of pathway enzymes. These compounds consist of a sugar core decorated with acyl groups, which are connected through ester linkages. In *Solanum lycopersicum* (tomato) four acylsugar acyltransferases (SlASAT1-4) sequentially add acyl chains to specific hydroxyl positions on a sucrose core leading to accumulation of tri and tetraacylated sucroses in the trichomes. To elucidate the spatial organization and interactions of tomato ASATs, we expressed SlASAT1-4 proteins fused with YFP in *N. benthamiana* and Arabidopsis protoplasts. Our findings revealed a distributed ASAT pathway with SlASAT1 and SlASAT3 localized to the mitochondria, SlASAT2 localized to the cytoplasm and nucleus, and SlASAT4 localized to the endoplasmic reticulum. To explore potential pairwise protein-protein interactions in acylsugar biosynthesis, we used various techniques, including co-immunoprecipitation, split luciferase assays, and bimolecular fluorescence complementation. These complementary approaches based on different interaction principles all demonstrated interactions among the different SlASAT pairs. Following transient expression of SlASAT1-4 in *N. benthamiana*, we were able to pull down a complex consisting of SlASAT1-4, which was confirmed through proteomics. Size exclusion chromatography of the SlASAT pulldown suggests a heteromultimeric complex consisting of SlASATs and perhaps other proteins involved in this interaction network. This study sheds light on the metabolic coordination for acylsugar biosynthesis through formation of an interaction network of four sequential steps coordinating efficient production of plant chemical defenses.

## Introduction

Plants produce a wide variety of structurally and functionally diverse metabolites. These compounds support plant growth, development, and adaptation to changing conditions, including biotic and abiotic stresses^1^. While advances in comparative transcriptomics and mass spectrometry have enabled the identification of entire plant specialized metabolic pathways^2^, the regulatory mechanisms and interactions between enzymes in these pathways are not well understood. One way in which metabolic pathways are regulated is through formation of protein-protein interaction networks. These multienzyme assemblies increase the local accumulation of metabolites to enable faster reaction rates, isolate intermediates from competing pathways, and protect the cell from reactive metabolites^3,4^. Metabolic complexes have been identified for some core and specialized metabolic pathways in plants and other organisms^4^ including diverse classes of specialized metabolites such as flavonoids, alkaloids, and cyanogenic glycosides^5–9^. A key feature in some of these metabolic complexes is scaffolding proteins that help stabilize the complex and allow coordination across subcellular compartments^6,10–12^.

Acylsugars are a class of specialized metabolites that play a crucial role in the intricate interactions between plants and their environment. The accumulation of acylsugars on glandular trichomes in the aerial parts of plants provide protection against biotic and abiotic stresses^13–18^. Acylsugars have a backbone sugar typically sucrose or glucose, with acyl groups of varying lengths, ranging from 2 to 16 carbon atoms, esterified at multiple hydroxyl positions^17,19^. *Solanum lycopersicum* (tomato) contains four trichome-localized BAHD family enzymes designated as SlASAT1 through SlASAT4, which are responsible for the biosynthesis of trichome-localized acylsugars^20–22^. SlASATs facilitate the esterification of acyl chains at specific positions on the sucrose backbone, leading to the production of mono-, di-, tri-, and tetra-acylated sucrose derivatives, however only tri and tetra acylsugars accumulate in planta^22^. Although, trichome-localized ASATs have been well studied in tomato and across the Solanaceae^23–25^, little is known about the localization and coordination of the acylsugar pathway. Here, we investigate potential metabolic complex formation in tomato trichome-localized acylsugar biosynthesis. Tomato acylsugar biosynthesis provides an excellent model to explore metabolic regulation and complex formation as it is localized to the trichomes, intermediates do not accumulate, and the pathway is enzymatically characterized. We find that SlASAT1 and SlASAT3 localized to the mitochondria, SlASAT2 showed dual localization in the cytosol and nucleus, and SlASAT4 was localized to the endoplasmic reticulum. Interestingly, despite the different organellar distribution of these ASATs, we found that they physically interact with each other using a variety of protein-protein interaction assays, including bimolecular fluorescence complementation, split luciferase, pulldown, co-immunoprecipitation, size exclusion chromatography, and proteomics. These findings suggest an acylsugar metabolic complex in tomato consisting of SlASAT1-4 and potentially other interacting proteins, with SlASAT2 acting as a key cytosolic link between other subcellular compartments. These findings are crucial for understanding the coordinated biosynthesis of acylsugars and enhancing crop resilience through accumulation of biologically active metabolites.

## RESULTS

### Tomato ASATs localize to distinct organelles

To experimentally determine the subcellular localization of tomato ASATs, we generated SlASAT fusion proteins with YFP and expressed them in *N. benthamiana* and Arabidopsis protoplasts. Prior to experimental determination of SlASAT1-4 subcellular localization, we predicted likely locations using subcellular prediction tools (Supplementary Fig. 1). In silico analyses provided an indication of subcellular localization, but the results were inconsistent for SlASATs across various prediction tools. SlASAT1 and SlASAT3 were predicted to localize to the mitochondria and cytosol (Supplementary Fig. 1). Similarly, SlASAT2 and SlASAT4 were predicted to localize mainly in the cytosol (Supplementary Fig. 1). The prediction guided placement of the YFP reporter gene, which was fused to the C-terminus of full-length SlASAT coding regions to minimize the chance of blocking a targeting signal. The resulting SlASAT-YFP constructs were transiently expressed in Arabidopsis protoplasts and *N. benthamiana* leaves. The corresponding YFP expression was analyzed using confocal microscopy. Localization in *N. benthamiana*, showed distinct patterns for the different SlASATs (Supplementary Fig. 2). SlASAT1 and SlASAT3 showed a punctate pattern that did not overlap with an endoplasmic reticulum (ER) marker protein or chlorophyll autofluorescence (Supplementary Fig. 2). SlASAT2 and SlASAT4 displayed a more continuous and distributed localization pattern that appeared to overlap with the ER marker protein, though cytosolic localization could not be ruled out (Supplementary Fig. 2). Furthermore, SlASAT2 exhibited a distinct localization, which appeared to overlap with the nucleus. SlASAT2 nuclear localization was confirmed using a nuclear-specific RFP marker (Supplementary Fig. 3).

Although, transient expression in *N. benthamiana* gave a sense for SlASATs localization, we used transfection in Arabidopsis protoplasts as a complementary approach and to provide more resolution to the subcellular localization. In Arabidopsis protoplasts expressing SlASAT1–YFP fusion, the YFP fluorescence was observed to localize to small punctate structures that co-localize with the mitochondrial marker (Fig. 1a), suggesting a mitochondrial localization. We further validated the localization of SlASAT1 by removing its predicted mitochondrial targeting signal (Fig. 1b and Supplementary Fig. 1b), which resulted in the enzyme localizing mainly to the cytosol and possibly the ER (Fig. 1b). Previous reports have shown that *N. benthamiana* ASAT1 is localized to the ER^26^. Thus, we also tested the SlASAT1-YFP localization with ER, golgi, and peroxisome marker proteins, and found that it did not overlap (Fig. 2 and Supplementary Fig. 4). This confirmed that SlASAT1 is indeed localized to the mitochondria (Fig. 1, 2a, and Supplementary Fig. 4a, c). In contrast, SlASAT2-YFP had a diffuse distribution, which did not overlap with the ER marker or chlorophyll autofluorescence, suggesting a cytosolic localization (Fig. 2b). To further validate the dual localization of SlASAT2, we co-expressed the SlASAT2-YFP fusion protein with a nuclear localization marker and observed the overlapping signal in the nucleus (Supplementary Fig. 3).

**Fig. 1.**
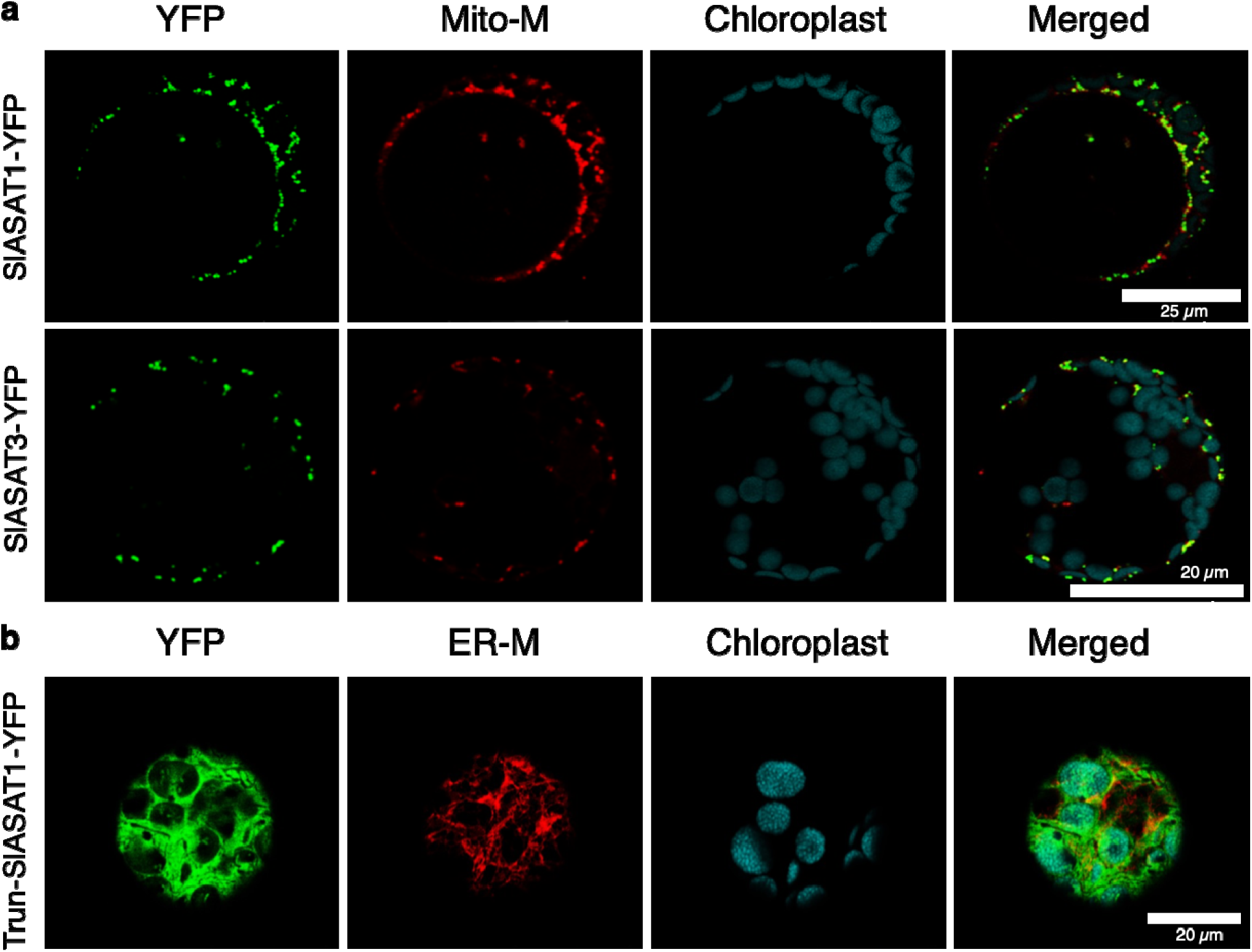
Subcellular localization of SlASAT1 and SlASAT3 fusion proteins in Arabidopsis protoplasts. **a** SlASAT1 and SlASAT3 fusion constructs (shown on the left) were expressed in Arabidopsis protoplasts and the localization of the corresponding protein was detected by confocal laser scanning microscopy. **b** SlASAT1-YFP without mitochondrial peptide fusion constructs (shown on the left, “Trun-SlASAT1-YFP”). The “YFP” panels (green) represent signals of SlASATs fused fluorescence proteins; the “Mito-M” panels (red) represent the signals of mitochondrial marker (MitoView™ Dyes); the “ER-M” panels (red) represent the signals of ER marker (AtWAK2-RFP-HDEL); the “chlorophyll” panels (cyan) represent chlorophyll autofluorescence; “Merge” panels show merged YFP, Mito-M, and chlorophyll signals. Scale bar shown in image. The experiments were repeated three times with similar results and representative images are shown.

**Fig. 2.**
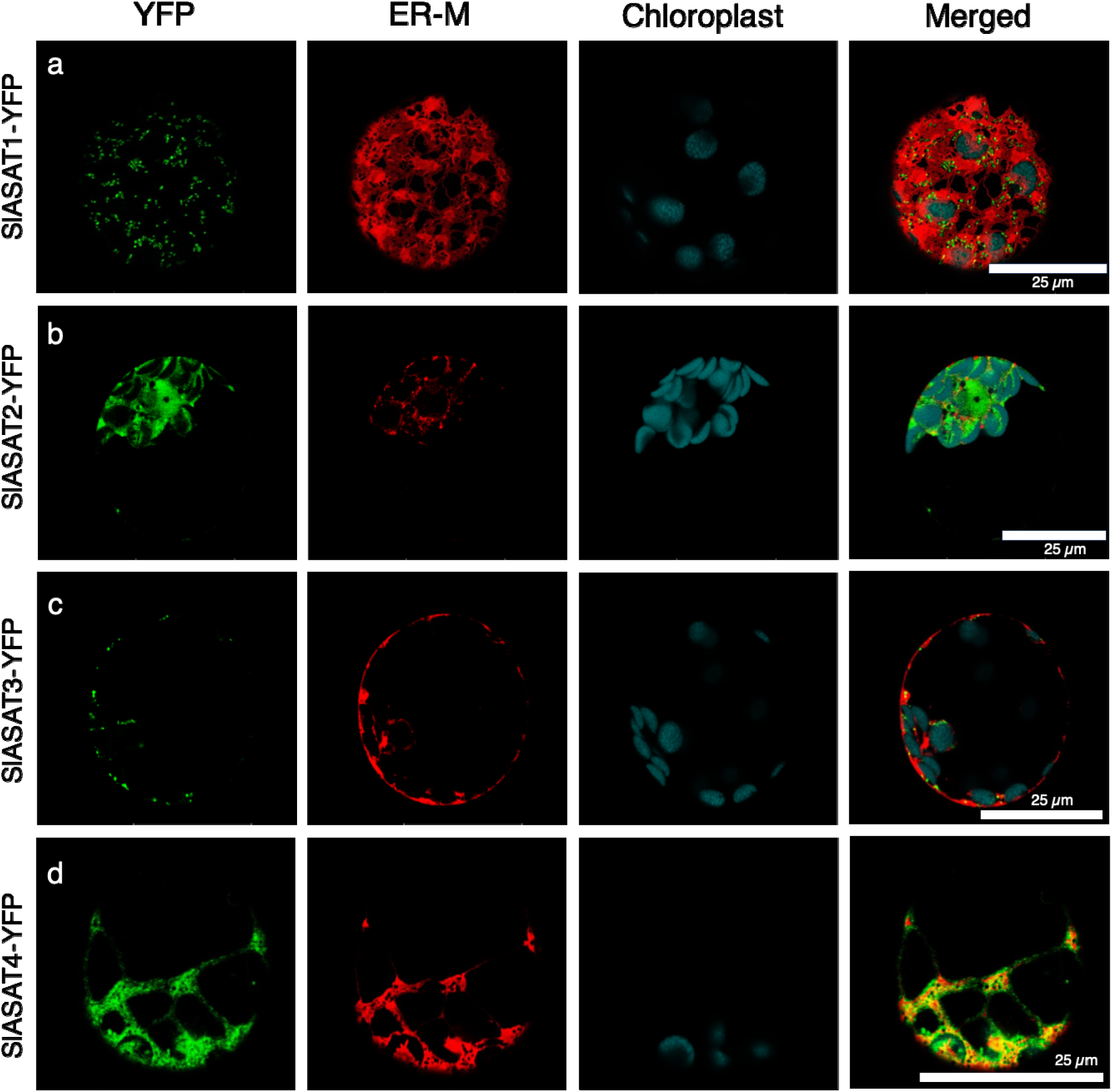
Subcellular localization of SlASAT1-4 fusion proteins in Arabidopsis protoplasts. SlASATs fusion constructs (shown on the left) were expressed in Arabidopsis protoplasts and visualized by confocal microscopy. The “YFP” panels (green) represent signals of SlASATs fused fluorescence proteins; the “ER-M” panels (red) represent the signals of ER marker (AtWAK2-RFP-HDEL); the “chlorophyll” panels (cyan) represent chlorophyll autofluorescence; “Merge” panels show merged YFP, ER-M, and chlorophyll signals. Scale bar shown in image. The experiments were repeated three times with similar results and representative images shown.

In silico predictions indicated that the SlASAT3 and SlASAT4 were localized to the mitochondria and cytosol, respectively (Supplementary Fig. 1a). These predictions were subsequently confirmed through transient expression in *N. benthamiana* and Arabidopsis protoplast transfection. The SlASAT3-YFP fusion protein exhibited a punctate subcellular localization pattern that co-localized with a mitochondrial marker, suggesting its mitochondrial localization (Fig. 1a). The localization of SlASAT3-YFP was further tested using markers for the ER (Fig. 2c), golgi, and peroxisomes (Supplementary Fig. 4b, d), and it was found that SlASAT3-YFP did not colocalize with any of these organelle markers, confirming its localization to the mitochondria. In contrast, SlASAT4 displayed a reticulon-like distribution that co-localized with an ER marker, indicating ER-specific localization (Fig. 2d and Supplementary Fig. 2d). Sequence analysis of SlASAT4 revealed the lack of a canonical ER-retention signal (KDEL/HDEL) in its amino acid sequence, suggesting an alternative ER-targeting mechanism^27^, potentially involving the N-terminal hydrophobic sequence. Subcellular localization studies demonstrate a distributed acylsugar pathway, whose coordination remained unknown.

### Co-transfection of SlASAT pairs alters subcellular localization

To determine the effect of binary protein-protein interactions on localization patterns, we infiltrated Arabidopsis protoplasts with combinations of SlASAT pairs one tagged with YFP and the other without. Plasmids expressing SlASAT1-YFP or SlASAT3-YFP were co-transfected with an untagged version of SlASAT2 into Arabidopsis protoplasts. Confocal microscopy revealed that co-infiltration of SlASAT1-YFP with SlASAT2 resulted in altered localization of SlASAT1 (Supplementary Fig. 5a). Interestingly, we found that SlASAT1 now localized to the cytosol (Supplementary Fig. 5a), instead of the mitochondria when expressed alone (Fig. 1a, 2a), suggesting that the presence SlASAT2 altered the localization of SlASAT1. We also tested the localization of SlASAT3-YFP in combination with SlASAT2, since SlASAT3 was also localized to the mitochondria (Fig.1a). Here, we observed a similar change in localization for SlASAT3, which localized to the cytosol (Supplementary Fig. 5b). These results suggest a potential role of cytosolic SlASAT2 in protein-protein interactions with other SlASATs.

### Complementary interaction techniques demonstrate SlASAT pairwise interactions

To test whether pairwise protein-protein interactions occur between SlASATs, we performed split luciferase complementation assays by expressing fusion protein pairs in *N. benthamiana* leaves. SlASAT1 and SlASAT3 were fused to the N-terminus of the luciferase fragment, while the potential interacting partners SlASAT2 and SlASAT4 were fused to the C-terminus. The assay indicated interactions between SlASAT1 and SlASAT2, SlASAT2 and SlASAT3, SlASAT3 and SlASAT4, and SlASAT1 and SlASAT4 (Fig. 3a and Supplementary Fig. 6). There was no evidence of an interaction in negative controls between nLUC-SlASAT1 and Empty vector (EV)-cLUC, or SlASAT4-cLUC and nLUC-EV, suggesting the observed SlASAT interactions are specific (Fig. 3a and Supplementary Fig. 6). We further quantified the luciferase assay using protein extracted from *N. benthamiana* infiltrated leaves and observed similar interactions among the SlASATs (Supplementary Fig. 6). Based on intensity of the relative luciferase units, the strongest interactions were observed for SlASAT1 and SlASAT2, and SlASAT3 and 4 (Supplementary Fig. 6b).

**Fig. 3.**
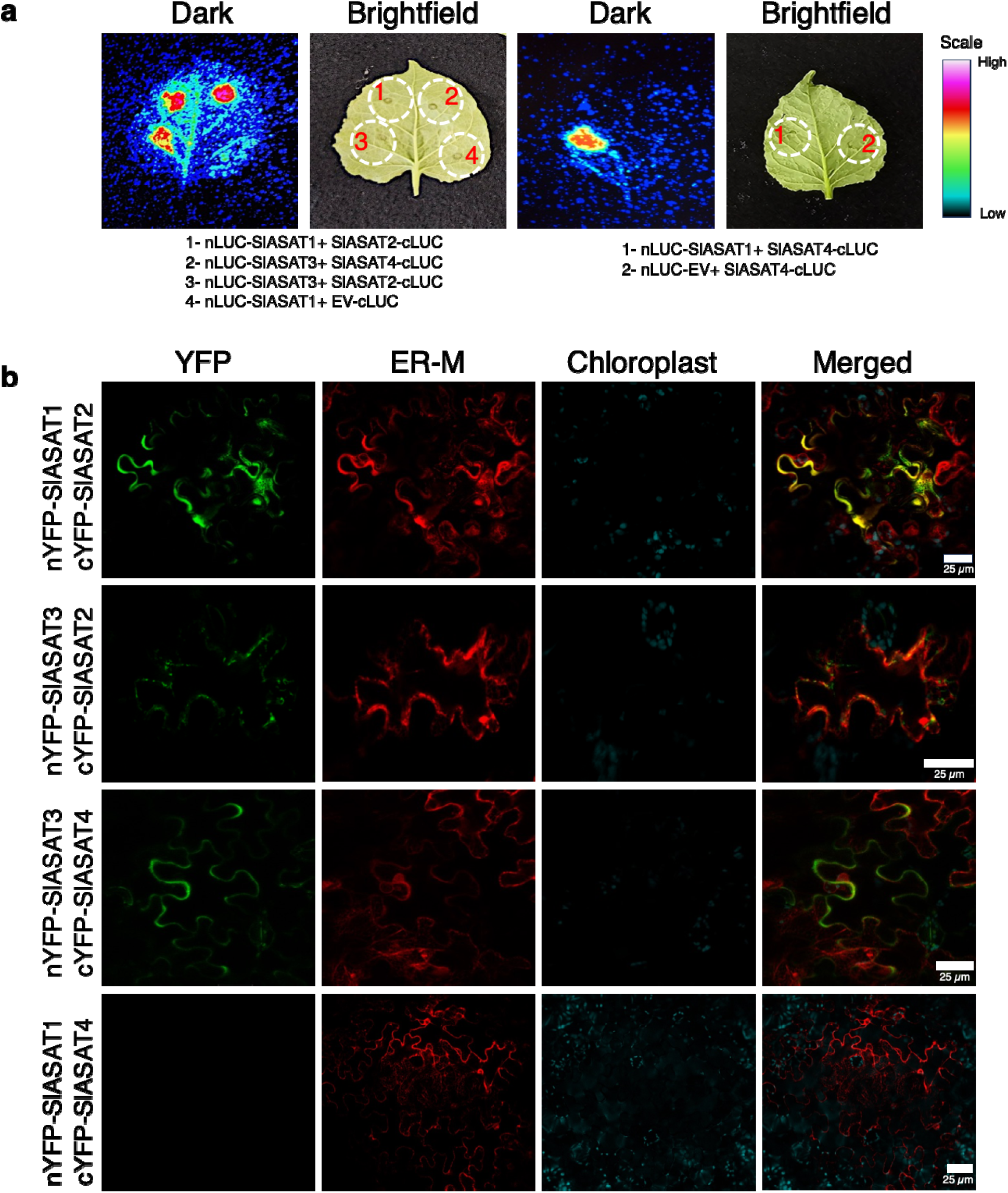
SlASAT protein-protein interactions via split-luciferase and bimolecular fluorescence complementation (BiFC) assay in *N. benthamiana*. **a** Split-luciferase assay image of *N. benthamiana* leaves co-infiltrated with *Agrobacterium* strains containing SlASATs following incubation for 3 days. Dotted circles indicate leaf panels that were infiltrated with *Agrobacterium* carrying the respective constructs. The luminescence intensity was recorded using a Photek-216. Scale bar indicates intensity of the observed luminescence. **b** BiFC assay of the interaction between SlASATs transiently expressed in *N. benthamiana* leaves. SlASAT1 and 3 were fused to N-terminal YFP, while SlASAT2 and 4 were fused to C-terminal YFP. The “YFP” panels (green) represent signals of SlASATs complex; the “ER-M” panels (red) represent the signals of ER marker (AtWAK2-RFP-HDEL); the “chlorophyll” panels (cyan) represent chlorophyll autofluorescence; “Merge” panels show merged YFP, ER-M, and chlorophyll signals. Scale bar shown in image. The experiments were repeated three times with similar results.

To further investigate pairwise SlASAT interactions, bimolecular fluorescence complementation (BiFC) assays were performed. The protein interactions were visualized using confocal microscopy, with the presence of fluorescence indicating either close proximity or interaction between the enzymes, as well as the subcellular localization of these interactions. The results showed that when SlASAT1 and SlASAT2 were used to complement YFP, strong fluorescence was observed that appeared to co-localize with the ER marker (Fig. 3b). SlASAT2 and SlASAT3 also displayed interaction, but to a lesser extent than SlASAT1 and SlASAT2. The interaction was detected in the cytosol, but it was difficult to determine whether these interactions occurred in the ER or the cytosol, based on the co-localization with the ER marker (Fig. 3b). SlASAT3 and SlASAT4 also displayed interactions (Fig. 3b). However, no interaction was observed between SlASAT1 and SlASAT4 (Fig. 3b), or between SlASAT2 and SlASAT4 (Supplementary Fig. 7), similar to negative controls that showed no complementation of the YFP signal (Supplementary Fig. 7). The level of fluorescence observed was indicative of the proximity of the interacting proteins, with stronger fluorescence suggesting more direct interactions and weaker fluorescence indicating associations through an intermediary or a complex of proteins. Consistent with split luciferase assays, strong interactions were detected between the SlASAT1/SlASAT2 and SlASAT3/SlASAT4 protein pairs (Fig. 3b), suggesting that these enzymes may directly interact with each other. Our results suggest that despite individual SlASATs being localized to different organelles, the acylsugar pathway enzymes can interact with each other, and these interactions may lead to changes in their subcellular localization as observed in the co-expression of SlASAT1/3-YFP with untagged SlASAT2 (Fig. 3b and Supplementary Fig. 5).

### Pairwise interactions of SlASATs via co-immunoprecipitation

To examine the interactions among the different SlASATs and further verify pairwise interactions, we performed co-immunoprecipitation (Co-IP) of tagged SlASATs transiently expressed in *N. benthamiana*. We first optimized the expression of the four SlASAT enzymes by individually expressing them in *N. benthamiana* and analyzing their expression levels in the total protein extracts and the immunoprecipitated fractions (Supplementary Fig. 8, 9). We further confirmed the western blot results through proteomics, which detected more than 55% sequence coverage of the SlASATs in the respective pulldowns (Supplementary Fig. 8, 9). Following optimization, we performed Co-IP experiments to investigate the interactions between the SlASATs.

As shown in Fig. 4a and c, SlASAT2 co-immunoprecipitated with HA-tagged SlASAT1. Additionally, SlASAT1 and SlASAT2 interacted in a reciprocal manner, with SlASAT1 being co-immunoprecipitated with Flag-tagged SlASAT2 (Fig. 4b and d). SlASAT4 co-immunoprecipitated with HA-tagged SlASAT3 (Fig. 4a and c) and the reciprocal interaction was observed through SlASAT3 being co-immunoprecipitated with Flag-tagged SlASAT4 (Fig. 4b and d). Similarly, SlASAT2 and SlASAT4 co-immunoprecipitated with the HA-tagged SlASAT3 and SlASAT1 (Supplementary Fig. 10). In contrast, no interaction was observed between Flag:RFP and SlASAT1 (Supplementary Fig. 10). The results further demonstrate pairwise interactions between SlASAT1 and SlASAT2, and SlASAT3 and SlASAT4 (Fig. 4a-d).

**Fig. 4.**
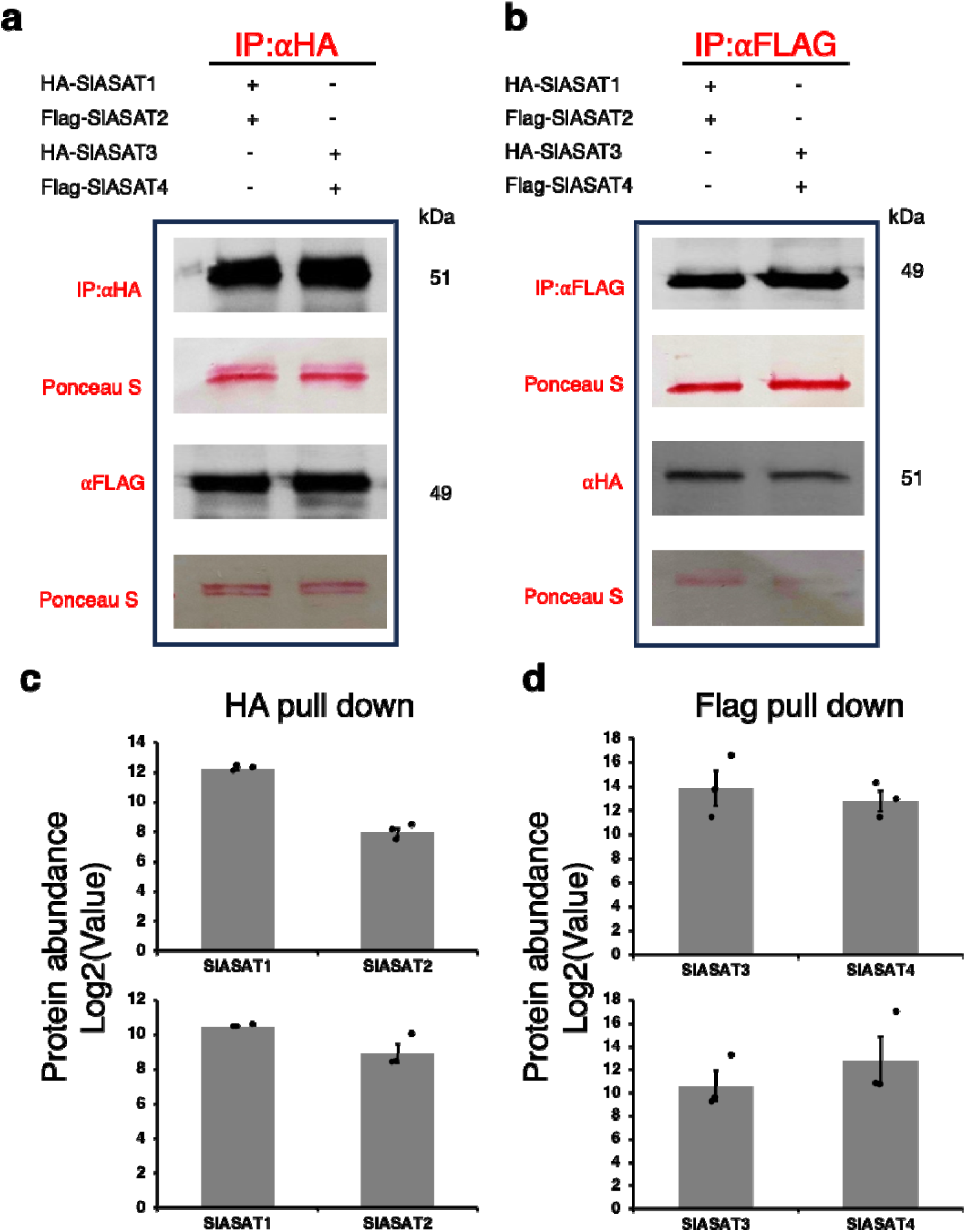
Co-immunoprecipitation of SlASATs show pairwise interactions. HA– and Flagtagged fusion proteins were transiently expressed in *N. benthamiana* leaves, and total proteins were extracted 3 days after infiltration. Protein extracts were transiently expressed in a pairwise manner: 35S::HA-SlASAT1 together with 35S::Flag-SlASAT2, and 35S::HA-SlASAT3 together with 35S::Flag-SlASAT4. **a** Protein gel blot analysis of HA-IP samples. IP was performed with anti-HA magnetic bead conjugate and interacting proteins were analyzed with an anti-Flag antibody. **b** Protein gel blot analysis of Flag-IP samples. IP was performed with anti-Flag magnetic bead conjugate, and interacting proteins were analyzed with an anti-HA antibody. **c**-**d** Quantification of the abundance of the HA/Flag pull-down metabolic complex using proteomics. Protein quantification was based on peptide spectral counts. Bars indicate average ± standard error of 3 biological replicates, with individual replicates plotted. Abbreviations: IP, Immunoprecipitation; HA, Hemagglutinin.

### SlASAT1-4 complex formation

Given the observed interactions between pairs of SlASATs, we next investigated if all four SlASATs formed a metabolic complex. We expressed constructs with untagged SlASAT2 and SlASAT3, along with HA-tagged SlASAT1 and Flag-tagged SlASAT4, in *N. benthamiana* and subsequently performed immunoprecipitation experiments using either HA or Flag magnetic beads (Fig. 5). The pulldown samples were initially confirmed by western blot (Supplementary Fig. 11) and then subjected to proteomic analysis. Our results showed that all four SlASATs were detected in the proteomics data of the pulldowns, indicating the formation of a multi-enzyme complex involved in acylsugar biosynthesis (Fig. 5b). In the Flag pulldown, SlASAT4 had greater abundance in this complex compared to SlASAT1, SlASAT2, and SlASAT3 (Fig. 5b). The differences in protein abundances observed may indicate the stoichiometry of the SlASAT complex, their expression levels in *N. benthamiana*, or their detectability in our proteomics analysis. These data indicate an acylsugar interaction network consisting of at least SlASAT1-4, representing four sequential steps of acylsugar biosynthesis. To gain insights into the molecular weight and stoichiometry of the SlASAT protein complex, we performed size exclusion chromatography (SEC) on the immunoprecipitated SlASAT complex eluted under non-denaturing conditions. Molecular weight standards for globular proteins were run prior to the SlASAT samples (Supplementary Fig. 12d). As shown in Fig. 5a, we observed three main protein peaks. These peaks correspond to a molecular weight of 271 kDa, 108 kDa, and 49 kDa, respectively (Fig. 5a). These data suggest a multimeric assembly consisting of SlASATs (approximate size of each SlASAT is 50 kDa). Western blot of the peak fractions with HA and Flag antibodies (targeting SlASAT1 and SlASAT4, respectively) demonstrate that SlASAT1 and SlASAT4 are found in the peak corresponding to the largest molecular weight, as well as other fractions (Supplementary Fig. 12). Together the SEC and western blot analysis suggests a heteromultimeric SlASAT complex (Fig. 5a and Supplementary Fig. 12).

**Fig. 5.**
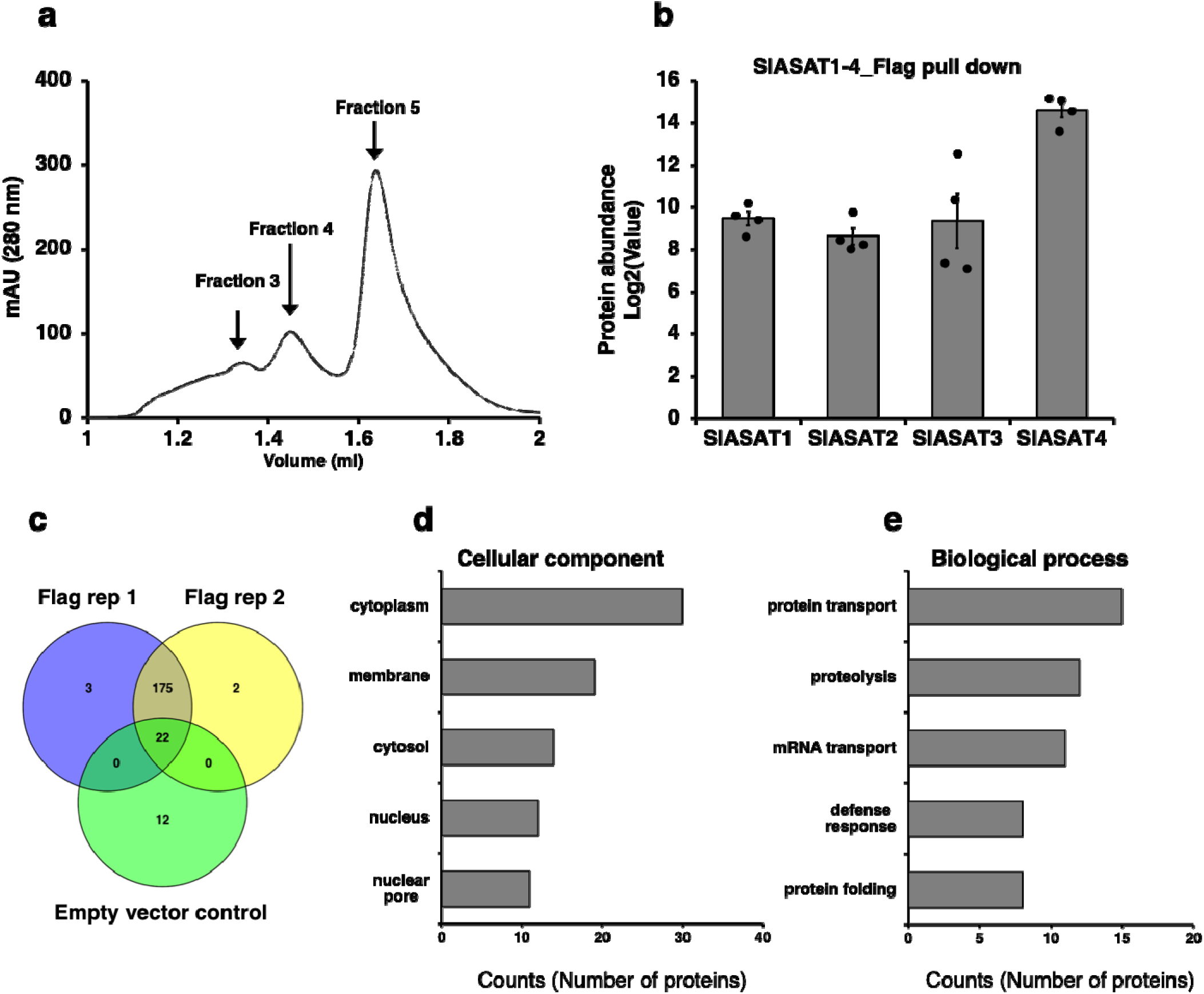
A multimeric SlASAT metabolic complex. HA-tagged SlASAT1 and Flag-tagged SlASAT4 were infiltrated in *N. benthamiana* together with untagged SlASAT2 and SlASAT3. Protein extracts were prepared from *N. benthamiana* leaves transiently expressing all four SlASATs and immunoprecipitation (IP) was performed using either anti-HA or Flag magnetic bead conjugates. **a** Size exclusion chromatography of the SlASAT1-4 complex. The SlASAT pulldown complex was eluted using non-denaturing conditions and separated using size exclusion chromatography. Peaks corresponding to ∼271 kDa (Fraction 3), ∼108 kDa (Fraction 4), and ∼49 kDa (Fraction 5) were identified and used for western blot analysis with HA and Flag antibodies (Supplementary Fig. 12). The molecular weight of the fraction was estimated based on protein standards run on an identical method and column (Supplementary Fig. 12). **b** Quantification of the abundance of the Flag pull-down metabolic complex using proteomics analysis. Protein quantification was based on peptide spectral counts. Bars indicate average ± standard error of 4 biological replicates, with individual replicates plotted. **c** Overlap of lists of interacting candidate proteins identified by LC-MS/MS analysis in two replicates and a negative (empty vector) control. **d** Functional enrichment analysis using QuickGo tool (https://www.ebi.ac.uk/QuickGO/annotations) of proteins Identified in the SlASAT1-4 metabolic complex pulldown experiment.

### Identification of additional interacting proteins in the SlASAT1-4 complex

Although our intent with Co-IP followed by proteomics was to characterize the SlASAT1-4 metabolic complex, we could also use this experimental setup to identify additional potential interacting proteins. To identify all proteins that might be interacting with the complex, we conducted a proteomics analysis of all proteins co-immunoprecipitated with the pull-down of SlASAT4 (Fig. 5c). Four biological replicates were prepared, with pull-down protein extracted using high detergent stringency for the SlASAT replicates and the EV controls. By mapping all detected proteins to the *N. benthamiana* proteome, we detected 175 unique proteins in the pull-down complex that were interacting with SlASATs, and we used the empty vector HA/Flag as a control to identify non-specific proteins (Fig. 5c). To identify potential true protein interactions, we hypothesized that those with abundant proteins detected in the pulldown are more likely to be true interactors. The relative abundance of the top 10 detected peptides in the pulldown experiments is shown in Supplementary Fig. 13 and Supplementary Table 2. The most abundant proteins detected in the pull-downs were involved in fatty acid biosynthesis, including acetyl-CoA C-acetyltransferase and Acyl-[acyl-carrier-protein] desaturase. Additionally, we identified transporter proteins such as ABC-type transporters and NbRanBP1-1a, which may play a role in shuttling intermediates in acylsugar biosynthesis^28–30^. Furthermore, we identified proteins involved in mitochondria-associated ER membrane contact sites. These proteins represent potential interactors with the ASAT acylsugar complex. To understand if the pulldown of the ASAT metabolic complex was enriched in proteins with other functions, we performed a functional enrichment analysis of the overlapping proteins in our pulldown replicates. Functional enrichment analysis for cellular process revealed enrichment of cytoplasm/cytosol and membrane among other terms (Fig. 5d). Functional enrichment for biological process indicated an enrichment of proteins related to transport, implying the diverse localization of the different complex components and potential interacting transporters in the ASAT complex.

## DISCUSSION

Since plants cannot rapidly evade harsh conditions, they rely on specialized metabolites to adapt and thrive in diverse environments. Although multi-enzyme complexes coordinating structurally diverse plant specialized metabolites have been characterized, for example in flavonoid biosynthesis, structural polymers, alkaloids, cyanogenic glycosides, and terpenoids^5–7,10,11,31–35^ it remains unclear how widespread a phenomena metabolic complexes are in regulating biosynthetic pathways. Here, we identified the multi-enzyme complex formation in tomato leading to acylsugar biosynthesis. We identified the key players of these interactions and, we show that despite SlASATs being localized to different organelles, they can interact with each other, adding another group of metabolites that are regulated via metabolic complex formation.

Tomato contains four trichome-enriched BAHD acyltransferases, SlASAT1-4, which sequentially acylate sucrose to produce tetra acylsucroses^20–22^. Despite the structural simplicity of acylsugar precursors the Solanaceae family exhibits remarkable acylsugar diversity^14,23^. This diversity is attributed to variations in the acylation pattern, branching pattern, and chain length (2-16 carbons) of the acyl groups. The SlASAT1-4 enzymes produce sequential acylation at the R4 (monoacylated), R3 (diacylated), R3’ (triacylated), and R2 (tetraacylated) positions, respectively, to generate acylated sucrose molecules^14,18,22,36^. The acyl-CoA substrates for each SlASAT are distributed across different organelles.

The upstream pathways that provide the acyl chains for ASATs are localized to various subcellular compartments^17,37^. The branched chain amino acids, which are the precursors to acyl chains, are biosynthesized in the plastids ^38^. Prior to esterification, the branched-chain amino acids are converted into their subsequent acyl-CoAs by the branched-chain α-keto acid dehydrogenase complex in the mitochondria^39^. These small CoAs are then esterified onto the sugar backbone by ASATs. Until now, ASATs were thought to be localized solely to the ER^26^, but here we demonstrate that SlASATs are not only found in the ER, but also in the mitochondria and cytosol (Figs. 1, 2, and Supplementary Figs. 2, 3, and 4). We show that SlASAT1 and SlASAT3 responsible for the first and third steps of this pathway, involving the acylation of short chain-CoAs (4 and 5 carbons), are localized within the mitochondria (Figs. 1a, 2a, c, and Supplementary Fig. 4) where they would have access to the short branched CoAs that are synthesized in the mitochondria. Short acyl-CoAs can undergo elongation likely in the plastids by a 2-carbon elongation mechanism^40,41^. SlASAT2 catalyzes acylation at the R3 position and is localized in the cytosol (Fig. 2b and Supplementary Fig. 3) and uses long (C10 and C12) acyl chains^22^. It is unclear how SlASAT2 accesses long acyl-CoAs in the cytosol. The final step, catalyzed by SlASAT4 uses acetyl-CoA as a substrate and acylates at the R2 position. Acetyl-CoA is typically synthetized in the plastids^39,42^, yet it is used as an intermediate in many metabolic processes and to post-translationally modify proteins, thus there may be a pool of acetyl-CoA available to SlASAT4 in the ER. The formation of an acylsugar metabolic complex may enable ASATs to access substrates in other compartments or through transfer from one SlASAT to another without intermediates being released.

Previous studies in diverse organisms have shown that membrane-bound organelles such as mitochondria, peroxisomes, chloroplasts, or the ER can physically interact coordinating various cellular processes, including metabolic pathways^43–45^. Reports of membrane-bound organelle interactions in plant metabolic pathways are limited, with the only known example being the glycolysis metabolic complex, which contains enzymes localized between the mitochondria and chloroplast that interact with each other^45^. However, in other systems, there are reports of ER mitochondrial contacts mediating fatty acid biosynthesis^43,44^. We speculate that similar membrane-bound organelle interactions may occur in tomato, given the differential localization of SlASATs. SlASAT2 may act as the inter-organellar linker as when SlASAT2 is expressed together with SlASAT1 or SlASAT3 it alters the localization of these enzymes (Supplementary Fig. 5).

We used various complementary experimental approaches, including BiFC, split luciferase assays, and Co-IP to confirm pairwise protein-protein interactions among SlASATs (Figs. 3, 4, and Supplementary Figs. 6, 7, 10). Additionally, we were able to pulldown the entire SlASAT complex and further validated these findings through proteomics and SEC analysis (Figs. 5 and Supplementary Fig. 11, 12). These results confirm the formation of a metabolic complex involving trichome-localized SlASATs. Tomato also makes acylsugars in the roots and the first step in the root-specific acylsugar pathway is catalyzed by a paralog of the trichome enriched SlASAT1^46^. It is possible that root acylsugar biosynthesis in tomato is also mediated by a metabolic complex consisting of root-specific SlASATs.

Metabolic complexes consist of the core metabolic enzymes, but often contain other proteins that are required for complex assembly4. Of the 175 proteins identified in our SlASAT pulldown, some likely represent true contributors to the SlASAT metabolic complex. The most abundant proteins were annotated to be involved in transport, fatty acid biosynthesis, and specialized metabolism (Supplementary Fig. 13, and Table 2). Interestingly, our pulldown experiments identified previously characterized membrane-bound, organelle-associated proteins, which could potentially serve as a tethering protein for the SlASAT metabolic complex (Supplementary Table 2). Our functional enrichment analysis revealed an abundance of proteins involved in biological processes such as protein transport (Fig. 5e), which could be responsible for the interorganellar transport of pathway intermediate that appears required for acylsugar biosynthesis. Similarly, we observed an enrichment of proteins associated with cytosolic and membrane-related functions (Fig. 5d). These proteins may directly or indirectly affect the metabolic complex (Fig. 5d, e). Further experiments are needed to follow-up on additional proteins that may be involved in the identified acylsugar metabolic complex.

We have demonstrated a metabolic complex for tomato trichome-localized acylsugar biosynthesis (Fig. 6). An ASAT complex may be conserved across the Solanaceae species that make acylsugars. However, each species has its own acylsugar biosynthetic differences, for example presence of additional enzymes, rearrangement of enzymatic order, use of different acyl chains and sugar cores, and many other differences that remain unknown^14,23,25,47^. Thus, if an ASAT metabolic complex is conserved across the Solanaceae, it may appear slightly different in a species-specific manner. Structural modeling and high-throughput proteomics approaches could be used to validate ASAT complex formation in other species.

**Fig. 6.**
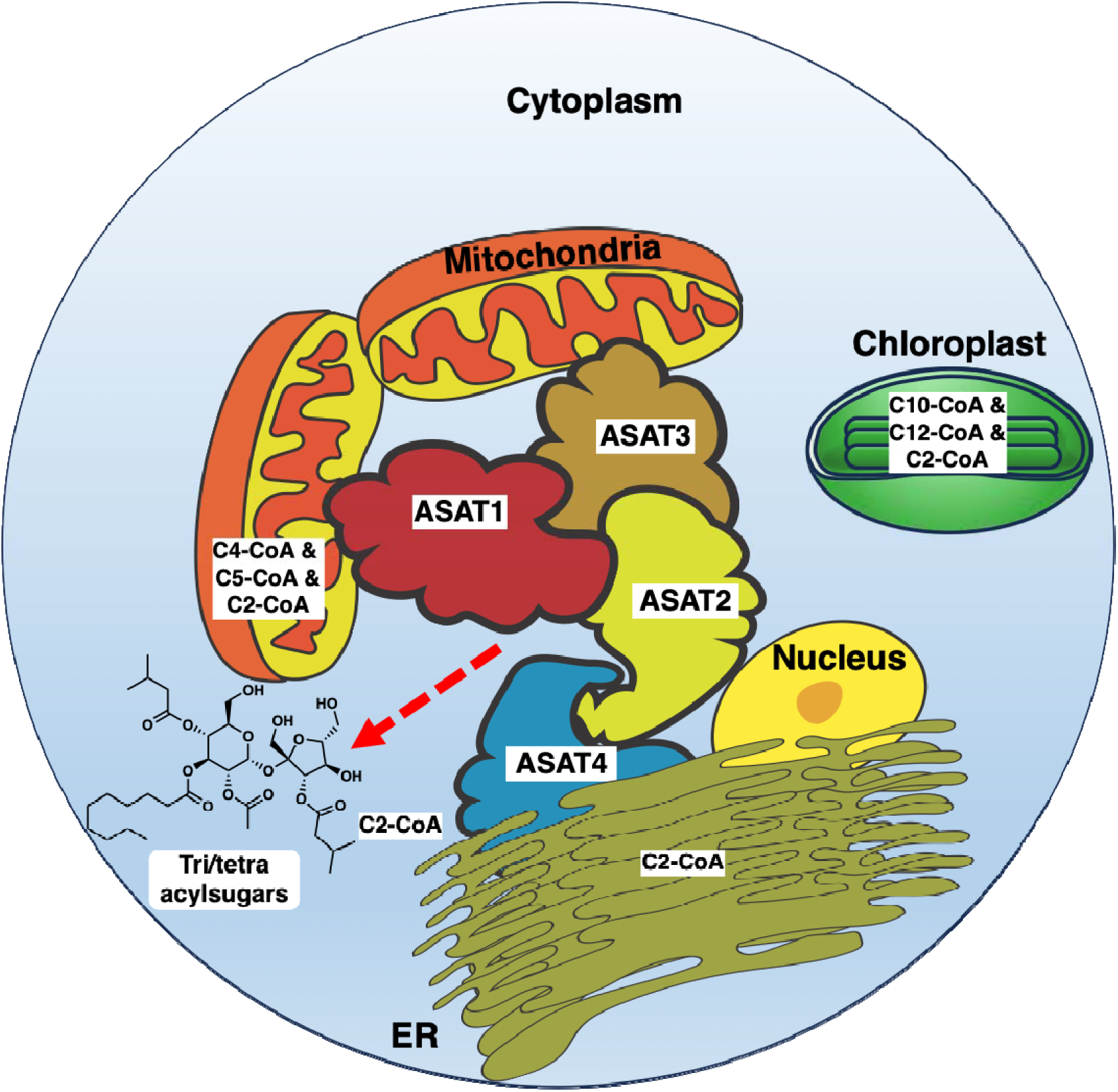
Proposed model for SlASAT metabolic complex. SlASATs are localized to different subcellular compartments yet form a metabolic complex. SlASAT2 is localized to the cytosol and may act as a key hub that links other ASATs localized to different compartments. Location of the pool of acyl chain CoA substrates is indicated. Red dashed line indicates release of tri and tetraacylsugars from the ASAT metabolic complex.

## Material and methods

### Subcellular localization of SlASATs

To determine the subcellular localization of SlASAT1-4, the full-length coding sequences for each gene were cloned using gene-specific primers (Supplementary Table 1) and transiently expressed in Arabidopsis protoplasts and *N. benthamiana* leaves. Briefly, the inserts were cloned into the pCAM-YFP plasmid using Gibson assembly, with the C-terminal enhanced YFP (eYFP) tag. For protoplast preparation^48^, Arabidopsis plants (ecotype Columbia) were grown under 16 h light at 23°C and 50% relative humidity. Leaves were thinly sliced with a razor blade and incubated with an enzyme solution containing 1% (w/v) cellulase (Goldbio), 0.25% (w/v) macerozyme (Goldbio), 0.4 M mannitol, 20 mM 2-Morpholinoethanesulfonic acid (MES) (pH 5.7), and 20 mM KCl incubated for 10 min at 55°C, and then 0.1% (w/v) bovine serum albumin and 10mM CaCl_2_ were added. The cut leaf tissues were incubated for 3 h at room temperature. The cell digest was filtered through 70-mm nylon mesh, and the protoplasts were harvested by centrifugation for 2 min at 100 × *g*, washed twice in 5 mL of ice-cold W5 solution (154 mM NaCl, 125 mM CaCl_2_, 5 mM KCl, 5 mM glucose, and 2 mM MES, pH 5.7), and then incubated with 2 ml of W5 solution for 30 min on ice. The pellet (protoplasts) was suspended in 1 mL of MMG solution (15 mM MgCl2, 0.4 M mannitol, and 4 mM MES, pH 5.7). Plasmid DNA (10 μg), either individually or combined with organelle-specific markers in a 1:1 ratio, was added to 100 μL of protoplast. Subsequently, 110 μL of a polyethylene glycol solution containing 40% polyethylene glycol-4000 (Sigma), 0.4 M mannitol, and 0.1 M CaCl_2_ was gently mixed with the protoplast-DNA mixture and incubated for 10-15 minutes at room temperature. The transfection reaction was stopped using 440 μL of W5 solution and then pelleted by centrifugation for 2 min at 100 × *g*. The pellet was resuspended in 500 μL of WI solution (20 mM KCl, 0.5 M mannitol, and 4 mM MES, pH 5.7) and incubated at 23 °C for 16-20 h. For subcellular localization in *N. benthamiana*, the constructs were transformed in *A. tumefaciens* GV3101. A single colony of each construct was used to inoculate a 5 mL LB media culture supplemented with 50 μg/mL kanamycin, 35 μg/mL rifampicin, and 50 μg/mL gentamycin. The bacterial cultures were incubated overnight at 28 °C with shaking at 200 rpm. The cultures were then centrifuged at 3,000 × *g* for 10 minutes, and the cell pellets were washed once with 5 mL of infiltration buffer (10mM MES pH 5.7, 10mM MgCl_2_, 200 μM acetosyringone) and incubated for 1 h in the dark before being used to infiltrate 3-week-old *N. benthamiana* leaves. For combinatorial infiltrations, the OD_600_ for each strain was set to 0.5. *N. benthamiana* leaves were coinfiltrated, in a 1:1 ratio, with an organelle marker and the desired construct, and visualized after 72 hours. The following organelle-specific markers were used for analysis: mitochondrial marker (cytochrome c oxidase IV-RFP), endoplasmic reticulum marker (AtWAK2-RFP-HDEL), peroxisomal marker (RFP-IHHPRELSRL), golgi marker (AtGT14 –RFP), and nuclear marker (NLS-(SV40)-RFP)^49–51^. Additionally, we used a mitochondrial tracking dye (MitoView™ 405, Biotium) to detect mitochondria. Confocal microscopy of Arabidopsis protoplasts or whole *N. benthamiana* leaves was performed using a Leica TCS SP8 inverted confocal microscope, with a 100x oil immersion and 20x dry objective. Fluorescence of each construct was recorded separately, and the images were merged to determine co-localization between marker protein or dye and SlASATs. The excitation wavelength and emission bandwidth recorded for each fluorescent protein as well as chlorophyll autofluorescence were optimized by the default presets in the Leica 2.6 software (Leica) and were as follows: eYFP (excitation 514 nm, emission 519–583 nm), RFP (excitation 561 nm, emission 590–630 nm), chlorophyll autofluorescence (excitation 633 nm, emission 652– 721 nm). For co-expression experiments using multiple SlASATs, we co-transformed SlASAT1-YFP and SlASAT2 (no fluorescent tag) together in a 1:1 molar ratio along with the organelle-specific marker using polyethylene glycol-mediated transformation in Arabidopsis protoplasts. To generate SlASAT1 without the mitochondrial targeting signal, we removed the predicted mitochondrial targeting signal (Supplementary Fig. 1) and cloned the desired gBlock into pCAM plasmid, then co-transformed it into the protoplasts.

### Bimolecular fluorescence complementation assay

SlASATs were cloned into the pCAM-nYFP and pCAM-cYFP plasmids, which incorporate the N-terminal (amino acids 1–174) and C-terminal (amino acids 175–239) segments of eYFP, respectively, upstream of the gene of interest. Specifically, SlASAT1 and SlASAT3 were cloned into the pCAM-nYFP construct, while SlASAT2 and SlASAT4 were cloned into the pCAM-cYFP construct. *N. benthamiana* leaves were then co-infiltrated with a 1:1:1 combination of *A. tumefaciens* harboring the ER-RFP marker, pCAM-nYFP, and pCAM-cYFP constructs as described above. Transient expression and YFP fluorescence were monitored 72 hours post infiltration using confocal microscopy, as done for the subcellular localization studies. Interactions were evaluated by testing all possible combinations in both directions. Additionally, each construct was assayed with an empty vector control to monitor any background fluorescence resulting from interactions between the gene of interest and YFP fragments. The bimolecular fluorescence complementation assay was performed using three biological replicates, with leaf infiltration done on separate plants.

### Split-luciferase complementation imaging assay

The plasmid pCAM-nLUC/cLUC was digested using *Nco*I to clone SlASAT1 and SlASAT3 into the downstream of pCAM-nLUC, and with *Swa*I to clone SlASAT2 and SlASAT4 into the upstream of pCAM-cLUC. The generated constructs were sequenced using sanger sequencing to verify the correct sequence. The nLUC and cLUC constructs harboring the respective genes of interest were transformed into *A. tumefaciens* and infiltrated into *N. benthamiana* as detailed above. For combinatorial infiltrations, the OD_600_ for each strain was set to 0.5. After 48-72 hours, the infiltrated leaves were then used for the LUC activity measurement. 0.25 mM Coelenterazine (Goldbio) was sprayed once onto the leaves, and they were kept in the dark for 7 minutes to allow chlorophyll luminescence to decay. The luminescence was then monitored using a Photek 216 (Photek, Ltd).

### In vivo pull-down experiments

To perform the in vivo pull-down of SlASAT1-4, the full-length coding sequences for each target were amplified by PCR from tomato genomic DNA using HA and Flag-incorporated primers listed in Supplementary Table 1. The DNA fragments were amplified using Phusion DNA Polymerase, PCR products purified using a PCR cleanup kit (Qiagen) and cloned into the pCAM plasmid. The constructs were then transformed into *A. tumefaciens* (GV3101) electrocompetent cells. The transformants were grown on LB plates containing 50 mg/ml kanamycin, 35 mg/ml rifampicin, and 50 mg/ml gentamicin at 28 °C. A single colony was used to inoculate 5 ml of LB medium supplemented with antibiotics and grown overnight at 28 °C. The cells were centrifuged at 3,000 g for 10 minutes, and the pellet was resuspended in 5 ml of infiltration medium (10mM MES pH 5.7, 10mM MgCl_2_, 200 μM acetosyringone) and centrifuged again. The pellet was resuspended in 5 ml of infiltration medium and incubated at room temperature for 2 hours. *A. tumefaciens* GV3101 containing the indicated constructs were incubated in infiltration buffer for 1 h in the dark and subsequently used to infiltrate 3-week-old leaves. For combinatorial infiltrations, the OD_600_ for each strain was set to 0.5. The leaves were harvested 3 days post-infiltration, frozen in liquid nitrogen, and stored at −80 °C. For the detection of pairwise SlASAT interactions proteins were extracted by homogenizing leaf tissue in a buffer containing 150 mM Tris-HCl (pH 7.5), 150 mM NaCl, 5 mM ethylenediamine tetraacetic acid (EDTA), 1 mM phenylmethylsulfonyl fluoride (PMSF), and 1% Nonidet P-40 (NP-40), along with a protease inhibitor cocktail (HaltTM Thermo Fisher Scientific). The homogenates were then clarified by centrifugation at 16,000 × g for 25 minutes at 4 °C, and the soluble proteins were incubated with anti-HA (Pierce™ Anti-HA Magnetic Beads) or anti-FLAG (Pierce™ Anti-DYKDDDDK Magnetic Agarose) magnetic beads for either 1 h at room temperature or overnight at 4 °C, following the manufacturer’s protocol. After removing the unbound proteins, the beads were washed three times with a buffer containing 10 mM Tris-HCl (pH 7.5), 150 mM NaCl, 0.5 mM EDTA, and 0.1% SDS. The proteins were recovered by resuspending the beads in Laemmli sample buffer (125 mM Tris-HCl (pH 6.8), 20.0 % SDS, and 4.0 % SDS), denaturing them at 100°C for 5 minutes. The proteins were separated by 12 % SDS-PAGE gel electrophoresis and detected by immunoblotting with monoclonal mouse anti-FLAG (Sigma) antibodies (1:1,500 dilution) or monoclonal mouse anti-HA (Invitrogen) antibodies (1:1,500 dilution) and goat anti-mouse IgG horseradish peroxidase conjugate (Invitrogen, 1:10,000 dilution) as a secondary antibody. Protein bands were visualized with chemiluminescence reagents (Millipore Chemo-HRP kit). To pull down the entire SlASAT1-4 complex, either HA-tagged SlASAT1 or Flag-tagged SlASAT4 was used as a bait, while SlASAT2 and SlASAT3 remained untagged. After immunodetection, Ponceau staining was used as a control for equal protein loading. Blotted membranes were incubated in Ponceau staining solution (Thermo Fisher Scientific) for 5 min, before excessive solution was washed away with distilled H2O.

### Proteomic analysis of the pulldown complex

The pulldown protein complexes were acetone precipitated overnight. The protein pellets were then washed twice with acetone. Proteins were denatured/reduced with 6 M urea, 2 M thiourea, and 5 mM DTT. Proteins were alkylated with 15 mM iodoacetamide (IAA). Protein were then digested with LyC at 1:50 (enzyme:protein) for 3 h at 37 °C and then digested with trypsin (1:50, enzyme:protein ratio) overnight at 37 °C. Digested peptides were purified by Evosep tips. Data were acquired on a Bruker timsTOF Pro2 connected to the Evosep-One system, and data were searched against *N. benthamiana* protein database concatenated with SlASAT1-4 proteins using Spectronaut version 19.3. For all proteome analyses, we used an EvoSep One liquid chromatography system ^52^ and analyzed the samples with a 44 minute gradient eluting the peptides (or Evosep One 30 SPD program). We used a 15 cm × 150 μm internal diameter column with 1.5 μm C18 beads (Bruker PepSep) and a 10 µm internal diameter zero dead volume electrospray emitter (Bruker Daltonik). Mobile phases A and B were 0.1% formic acid in water and 0.1% formic acid in acetonitrile, respectively. Evosep One was coupled online to a modified trapped ion mobility spectrometry quadrupole time-of-flight mass spectrometer (timsTOF Pro 2, Bruker Daltonik GmbH, Germany) via a nanoelectrospray ion source (Captive spray, Bruker Daltonik GmbH). To conduct data-independent acquisition-parallel accumulation serial fragmentation (DIA-PASEF), the timsTOF Pro2 operated in long gradient DIA-PASEF mode. 16 scans with narrow 25 *m/z* isolation windows (resulting in 32 windows) covered an m/z range of 400 to 1200 and an ion mobility range of 0.6 to 1.43 Vs.cm−2. The duty cycle was maintained at 100%. This setup results in a total cycle time of 1.8 seconds (1x 100 ms MS1 survey scan, 16x 100 ms DIA-PASEF scans).

### Analysis of the Proteomics Data

The DIA-PASEF raw data was analyzed using the directDIA workflow of Spectronaut version 19.3 with default settings: Trypsin digestion with 2 allowed missed cleavages, cysteine carbamidomethylation as a fixed modification, methionine oxidation, and acetylation at the protein N-terminus as variable modifications. All searches were conducted against Uniprot *N. benthamiana* database concatenated with SlASAT1-4. Proteins and peptides were filtered at Q-value of 0.01. Functional enrichment analysis was performed using the QuickGO annotation tool (https://www.ebi.ac.uk/QuickGO/annotations) to classify the proteins into three categories: biological processes, cellular compartments, and molecular functions. For any identified proteins that were not annotated in the UniProt database, the InterProScan software was utilized to annotate their Gene Ontology functions.

### Size Exclusion Chromatography of ASAT1-4 complex

Following expression of SlASAT1-4 in *N. benthamiana*, the protein complex was pulled down using FLAG-tagged ASAT4 or HA-tagged ASAT1 as bait. Since the proteins were analyzed as complexes, elution was conducted using non-denaturing conditions with HA peptide. Samples were subsequently separated under native conditions by size exclusion chromatography using a Superdex 200 3.2/300 column (Cytiva) and equilibrated with 50 mM Tris-HCl, 150 mM NaCl, pH 7.4 at a flowrate of 0.050 mL/min. Prior to loading, samples were concentrated using 30 kDa molecular weight cutoff centrifugal filter units (Millipore, Amicon Ultra Centrifugal Filter). 50 µL of the sample was injected onto the column and elution was monitored by UV absorbance at 280 nm, and fractions were collected at 100 µL intervals. Molecular weight determination was achieved by comparison to gel filtration standard protein mixture (BioRad) monitored at UV absorbance at 215 nm under same elution conditions.

## AUTHOR CONTRIBUTIONS

VD conducted experiments, analyzed data, wrote and edited the manuscript. EO and AY conducted experiments, analyzed data and edited the manuscript. CAS conceived experimental setup, analyzed data, wrote and edited the manuscript.

## ACKNOWLEDGMENTS

We thank Alexander Jurkevich and Frank Baker from AMLC for assistance in confocal microscopy. We thank Brian Mooney and Thao Nguyen from the Charles W. Gehrke Proteomics Facility at MU for assistance in proteomics analyses. Research reported in this publication was supported in part by a The National Institute of General Medical Sciences grant awarded to AY [R35 GM155253].

