## Supplementary material for "A Metabolic Complex Involved in Tomato Specialized Metabolism": Supplememtary Figures 1-13

**a**

| Tools | SlASAT1 | SlASAT2 | SlASAT3 | SlASAT4 |
|---|---|---|---|---|
| **TargetP** (https://services.healthtech.dtu.dk/services/TargetP-2.0/) | Mitochondria | Other | Mitochondria | Others |
| **WoLF PSORT** ( https://wolfpsort.hgc.jp/) | chloroplast | chloroplast | Mitochondria | Cytoplasm |
| **Predotar** (https://urgi.versailles.inra.fr/predotar/) | Mitochondria | Possibly Mitochondria | Possibly Mitochondria | None |
| **Plant-mPLoc** (http://www.csbio.sjtu.edu.cn/bioinf/plant-multi/) | Cytoplasm | Cytoplasm | Cytoplasm | Cytoplasm |
| **Deeploc-2.0** (https://services.healthtech.dtu.dk/services/DeepLoc-2.0/) | Nucleus | Nucleus | Nucleus | Nucleus |

**b**

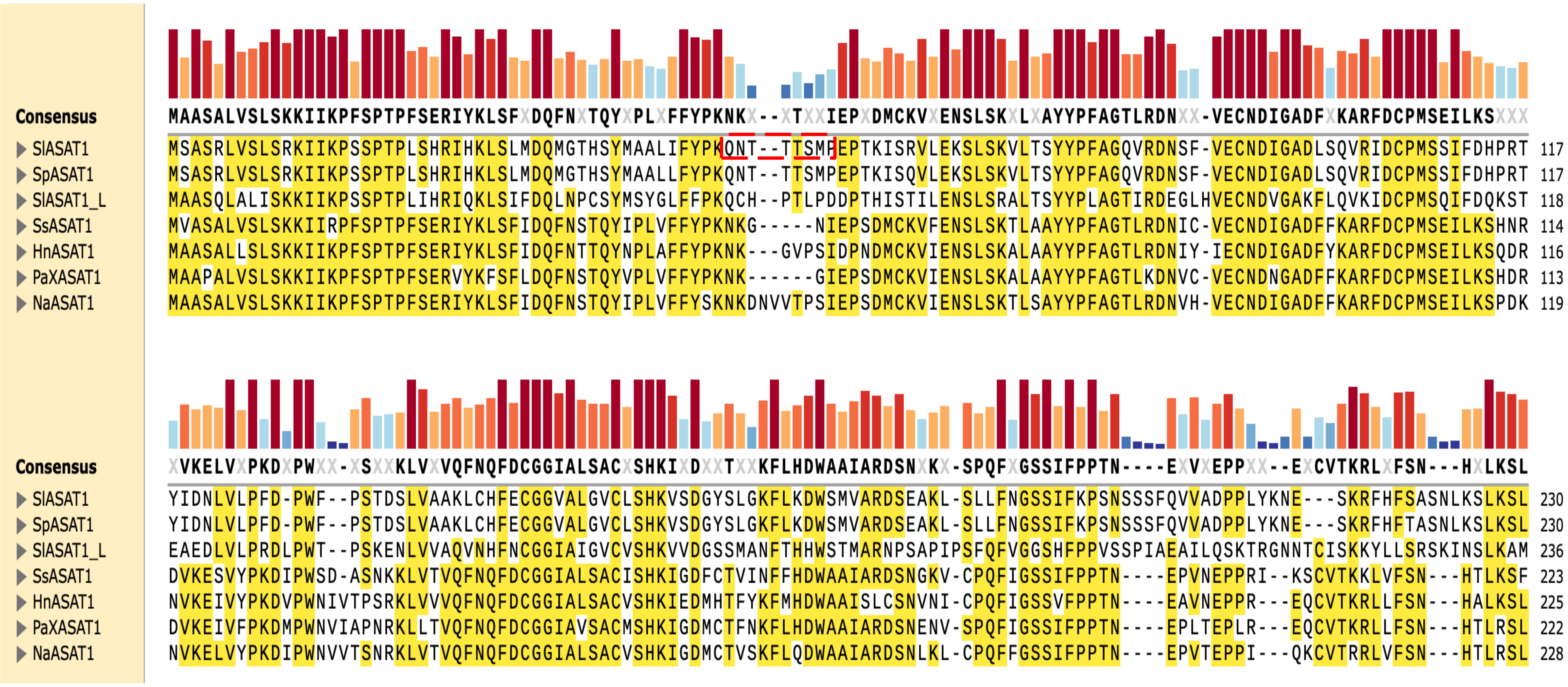


**Supplementary Fig. 1 In silico subcellular localization prediction of tomato ASATs using different tools. a** The subcellular localization was predicted using several tools, with the results indicated by the organelle scores. The scores represents the reliability of the prediction, with the organelle with the highest score shown for each tool. **b** The sequence alignment of previously characterized ASAT1. The sequence of SlASAT1 shows the predicted mitochondrial targeting signal highlighted in red box. SlASAT1 *Solanum lycopersicum*, SpASAT1 *Solanum pennellii*, SlASAT1-L *Solanum lycopersicum*-Like, SsASAT1 *Salpiglossis sinuata*, HnASAT1 *Hyoscyamus niger*, PaASAT1 *Petunia axillaris* , NaASAT1 *Nicotiana attenuata*. The bars above the alignment represent sequence identity, with red indicating 100%, orange (60-75%), yellow (50-60%), light blue (25-50%), and dark blue (< 20%).

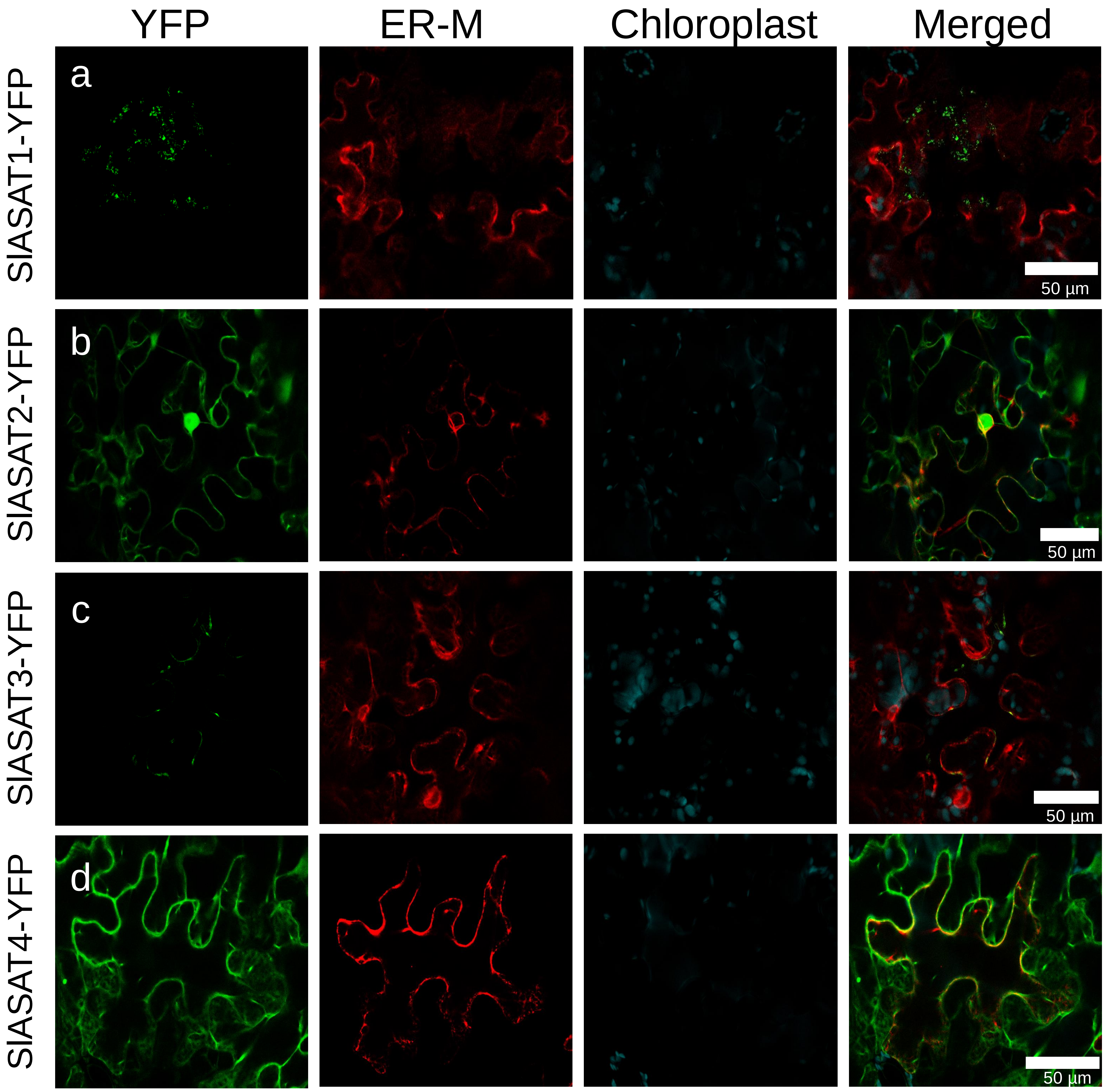


**Supplementary Fig. 2 Subcellular localization of SlASATs in *N. benthamiana* leaves.** SlASAT1-4-YFP fusion constructs were expressed in *N. benthamiana* leaves and the localization of the corresponding proteins were detected by confocal laser scanning microscopy 3 days after infiltration. The "YFP" panels (green) represent signals of SlASAT1, SlASAT2, SlASAT3, and SlASAT4 fused fluorescence proteins; the "chlorophyll" panels (cyan) represent chlorophyll autofluorescence; the "ER-M" panels (red) represent the signals of ER marker (AtWAK2- RFP-HDEL)*;* "Merge" panels show merged YFP, chlorophyll, and Marker. Representative images are shown with a scale bar shown in each image.

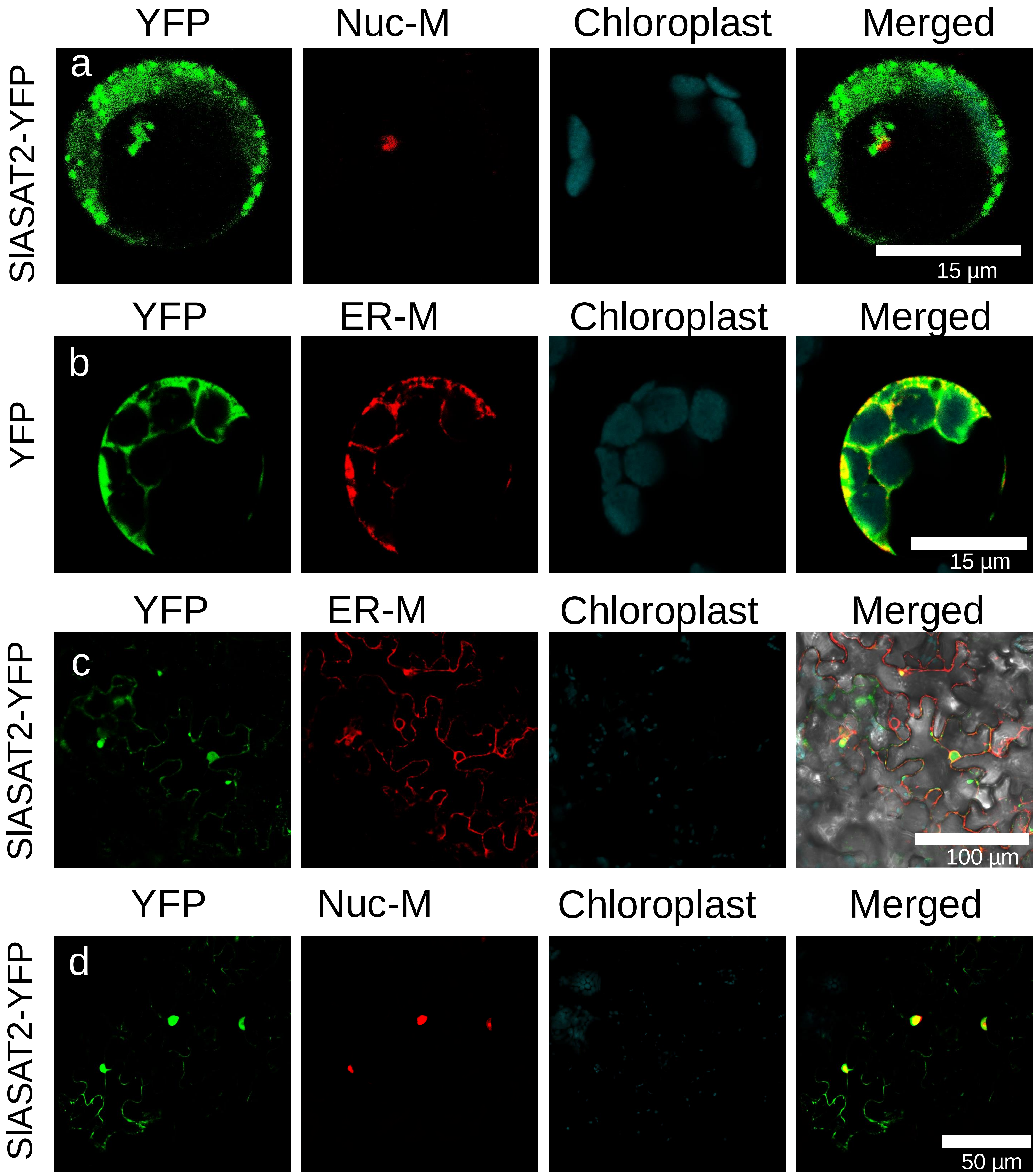


**Supplementary Fig. 3 SlASAT2 co-localizes with the nuclear marker in both Arabidopsis protoplasts and *N. benthamiana* leaves.** SlASAT2 fusion construct or free YFP (shown on the left) were expressed in Arabidopsis protoplasts (**a**-**b**) and *N. benthamiana* leaves (**c**-**d**) and the localization of corresponding proteins was detected by confocal laser scanning microscopy. The “YFP” panels (green) represent signals of SlASAT2 fused fluorescence proteins; the “NUC-M” panels (red) represent the signals of nuclear marker (SV40-RFP); the “ER-M” panels (red) represent the signals of ER marker (AtWAK2-RFP-HDEL); the “chlorophyll” panels (cyan) represent chlorophyll autofluorescence; “Merge” panels show merged YFP, organelle marker, and chlorophyll signals. Representative images are shown with a scale bar shown in each image.

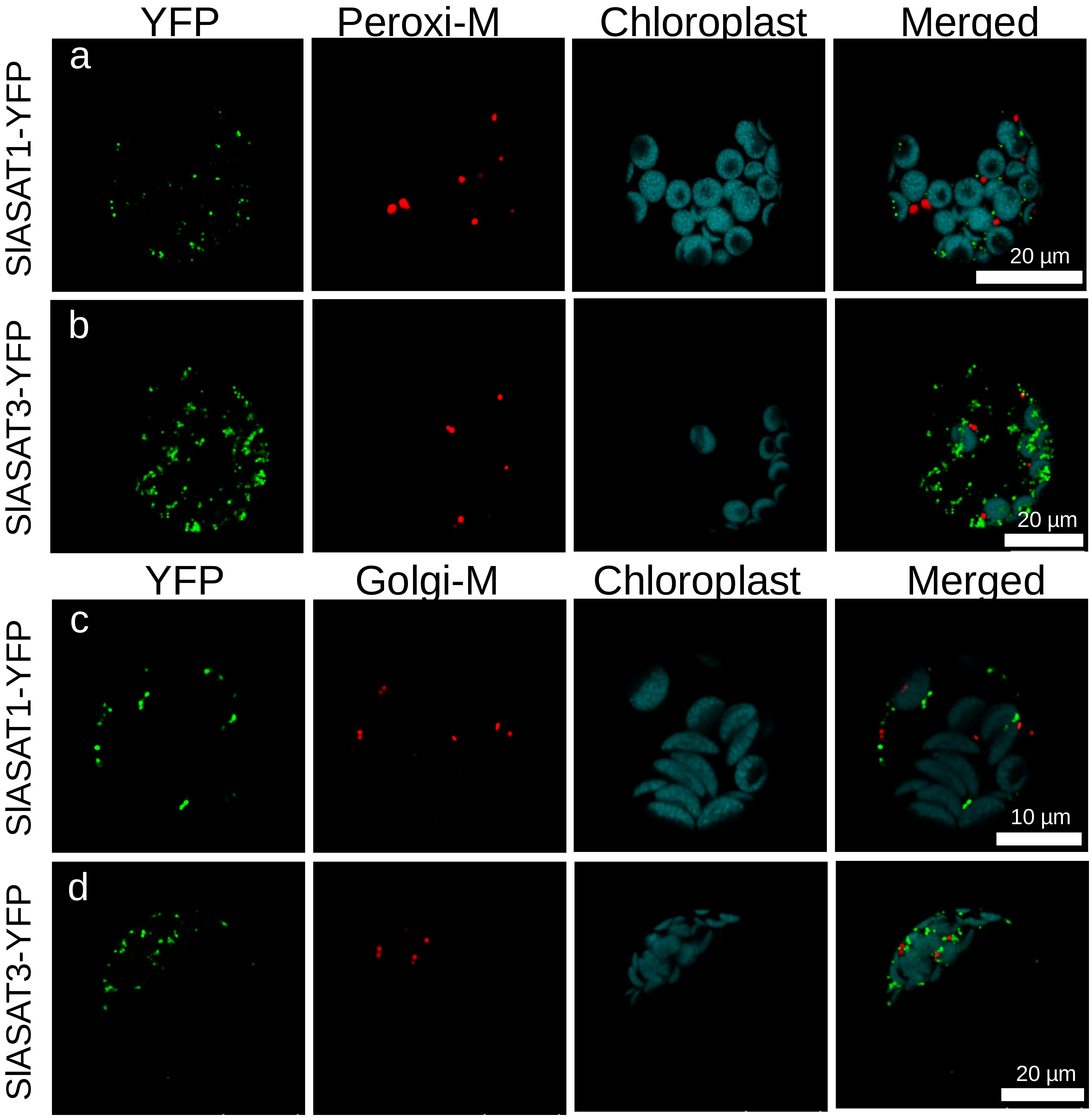


**Supplementary Fig. 4 Subcellular localization of SlASAT1 and SlASAT3 fusion proteins in Arabidopsis protoplasts.** The SlASAT1-YFP and SlASAT3-YFP fusion constructs (shown on the left) were expressed in Arabidopsis protoplasts and the localization of corresponding proteins was detected by confocal laser scanning microscopy. Panels **a**-**b** depict co-expression of SlASAT1 and SlASAT3 with a peroxisome marker, while panels **c**-**d** show co-expression of SlASAT1 and SlASAT3 with a Golgi marker. The "YFP" panels (green) represent signals of SlASATs fused fluorescence proteins; the "Peroxi-M" panels (red) represent the signals of peroxisome marker (RFP-IHHPRELSRL); the "Golgi-M" panels (red) represent the signals of golgi marker (AtGT14-RFP); the "chlorophyll" panels (cyan) represent chlorophyll autofluorescence signal; "Merge" panels show merged YFP, organelle marker, and chlorophyll signals. Representative images are shown with a scale bar shown in each image.

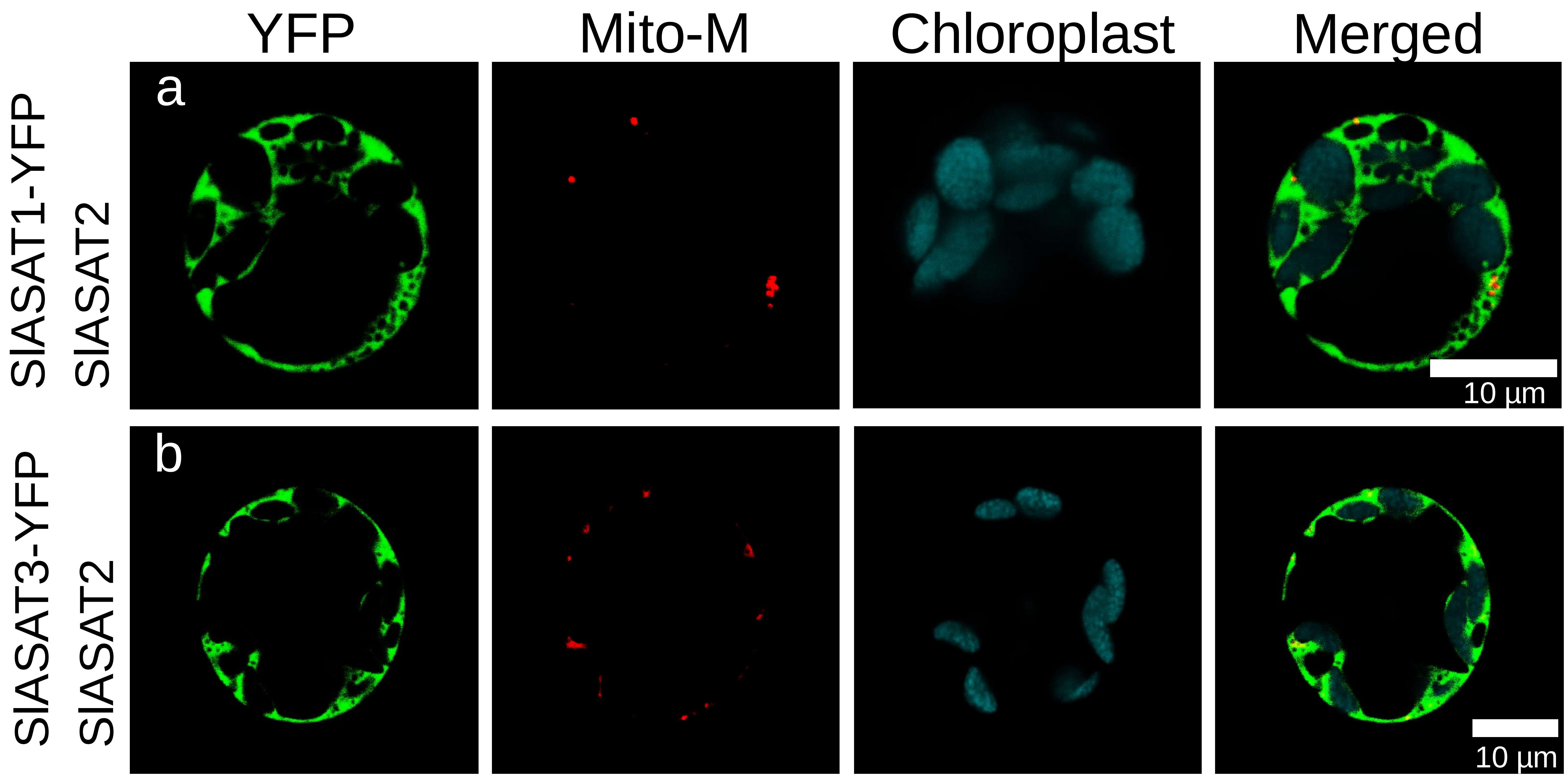


**Supplementary Fig. 5 Co-expression of SlASAT protein pairs alters their subcellular localization.** Co-expression of SlASAT1-YFP (**a**) and SlASAT3-YFP (**b**) together with untagged SlASAT2 changed their subcellular localization, preventing them from co-localizing with the mitochondrial marker. The "YFP" panels (green) represent signals of SlASATs fused fluorescence proteins; the "Mito-M" panels (red) represent the signals of mitochondrial marker (MitoView™ Dyes); the "chlorophyll" panels (cyan) represent chlorophyll autofluorescence; "Merge" panels show merged YFP, Mito-M, and chlorophyll signals. Representative images are shown with a scale bar shown in each image.

a

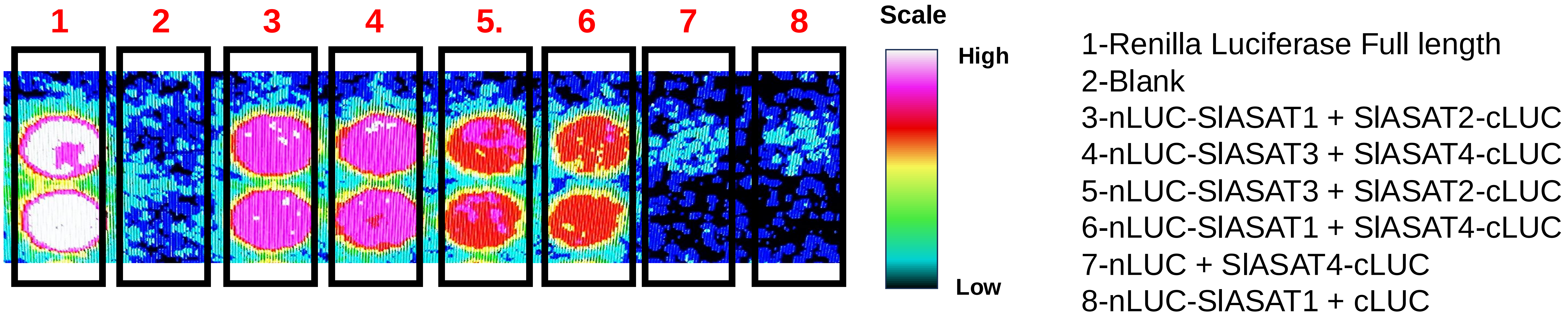


b

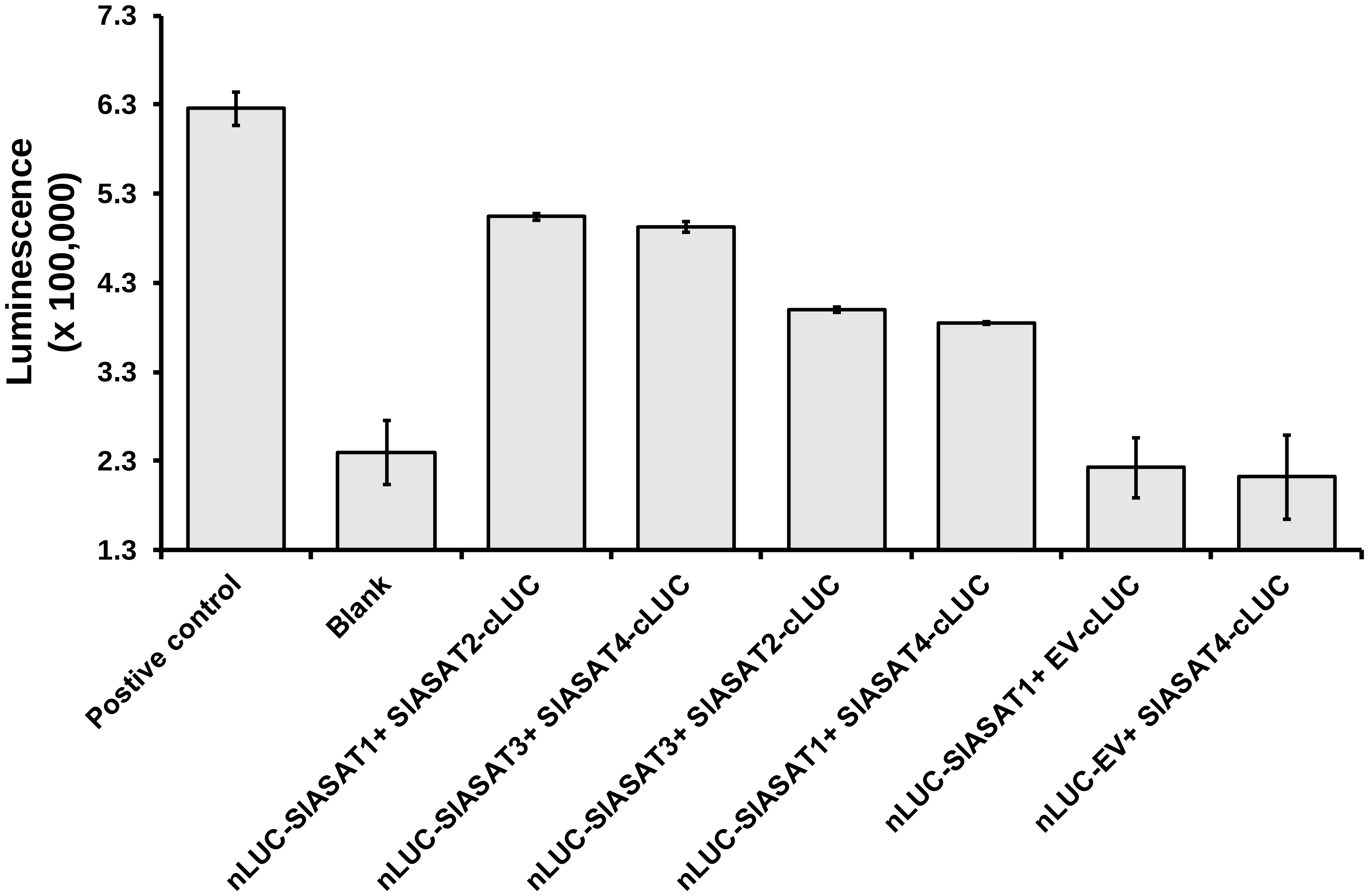


**Supplementary Fig. 6 Split-luciferase complementation assay in *N. benthamiana*.** (**a**) Split-luciferase assay images of protein extracts from *N. benthamiana* leaves expressing different SlASAT combinations. Proteins were extracted and placed in a 96 well plate and the luciferase activity was monitored in each well plate. The two replicate wells are shown for each construct. (**b**) Quantification of the relative luminescence unit intensity for different combination of SlASATs using Image J. Bars indicate average ± standard error of 2 biological replicates.

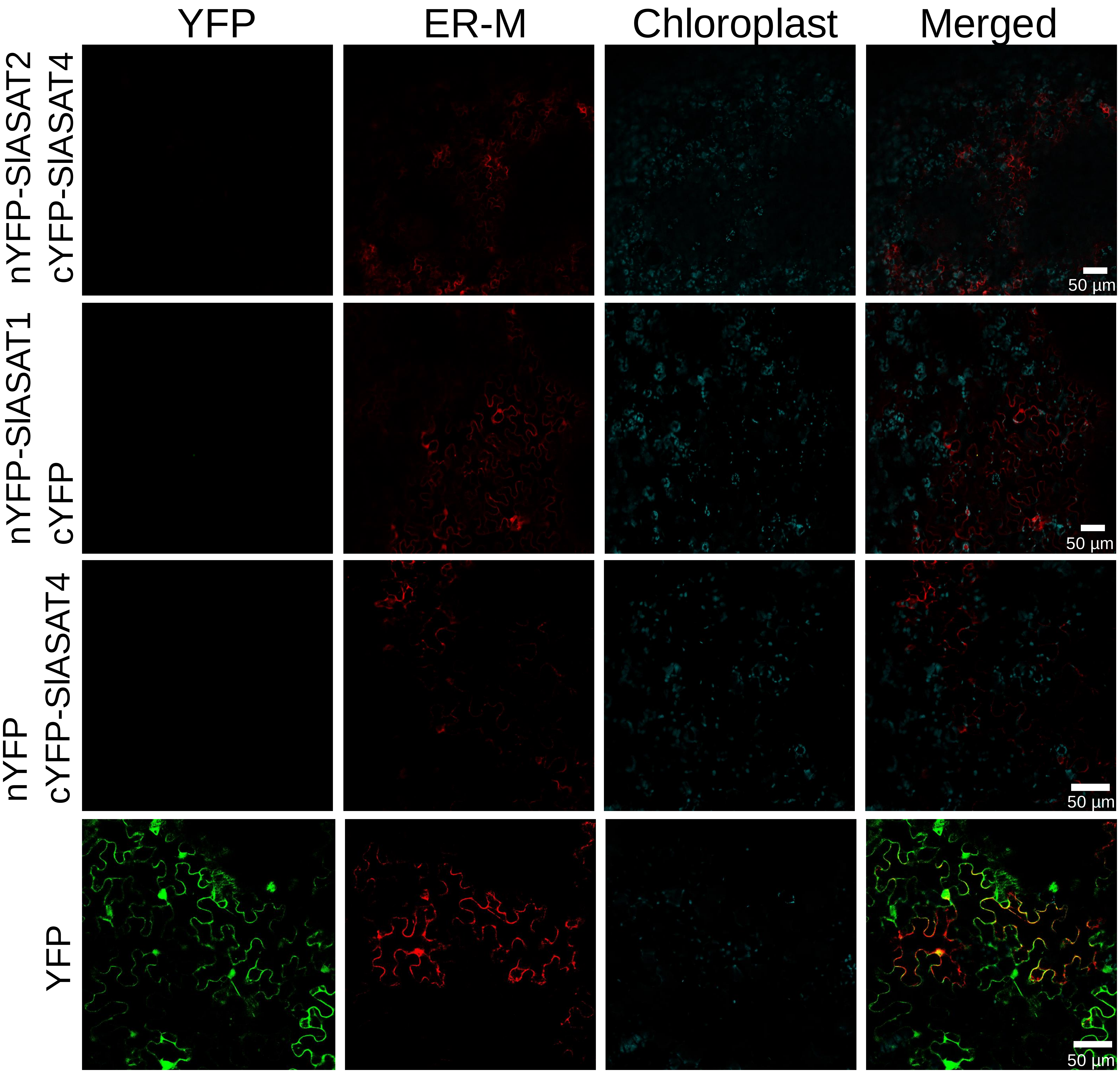


**Supplementary Fig. 7 SlASAT protein-protein interactions by BiFC in *N. benthamiana.*** BiFC assay of the interaction between SlASATs transiently expressed in *N. benthamiana* leaves. SlASAT1 and 3 were fused to N-terminal YFP, while SlASAT2 and 4 were fused to C-terminal YFP. Pairs of SlASAT1 +cYFP, and SlASAT4 +nYFP, were used as negative controls. The "YFP" panels (green) represent reconstituted YFP fluorescence; the "ER-M" panels (red) represent the signals of ER marker (AtWAK2-RFP-HDEL); the "chlorophyll" panels (cyan) represent chlorophyll autofluorescence; "Merge" panels show merged YFP, ER-M, and chlorophyll signals. Representative images are shown with a scale bar shown in each image.

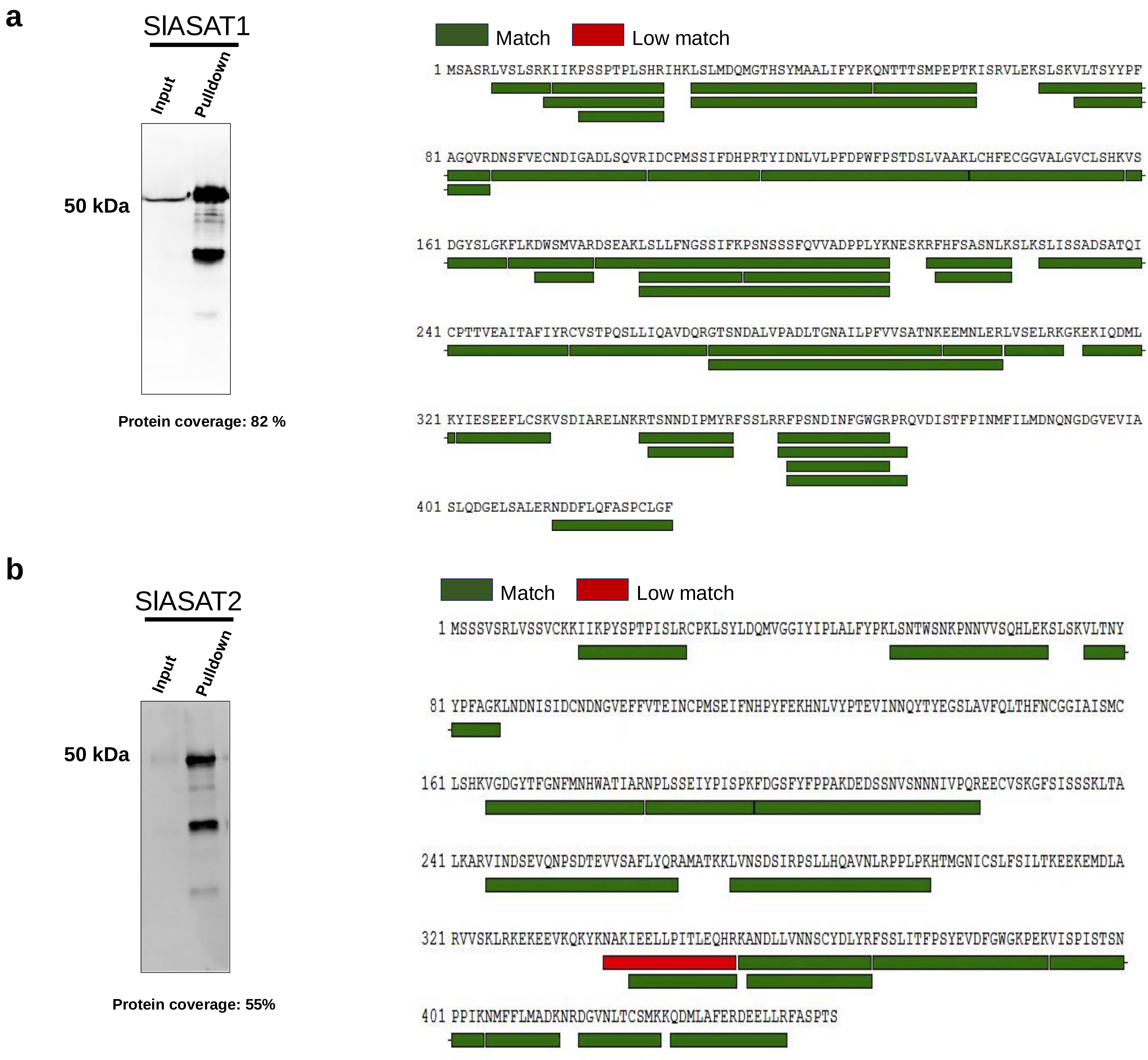


**Supplementary Fig. 8 SlASAT1 and SlASAT2 pulldown optimization.** Protein gel blot and proteomics analysis of SlASATs pulldown. The protein gel blot analysis of the input and HA-SlASAT1 (**a**) and Flag-SlASAT2 (**b**) immunoprecipitation (IP) samples, along with the schematic overview of the SlASAT proteins detected using LC/MS-MS analysis. IP was performed with anti-HA or anti-Flag antibody. The experiment was repeated at least three times, with similar results. The peptides identified in the LC/MS-MS analysis are represented by green boxes (high match > 90% amino acid identity), while the low match peptides are represented by red boxes (< 90% amino acid identity).

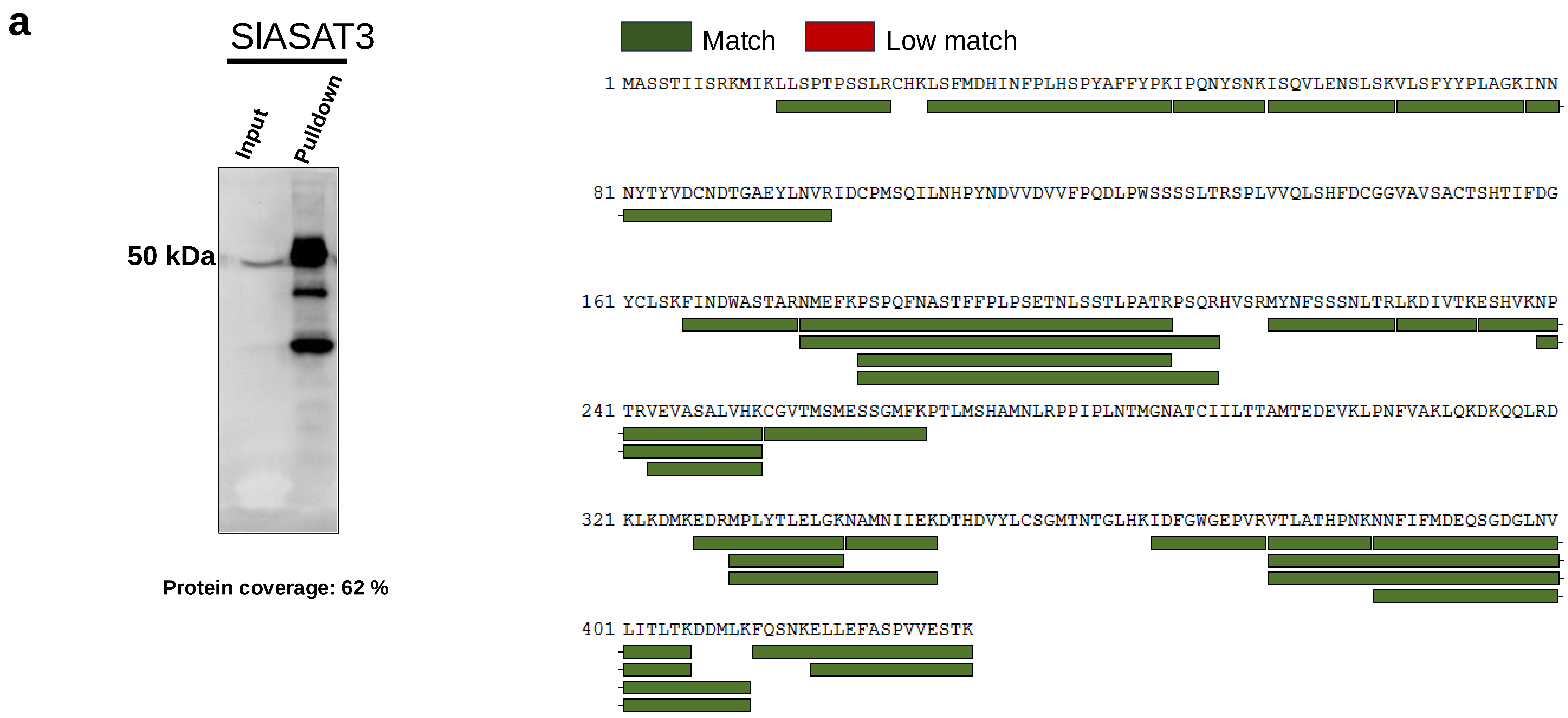


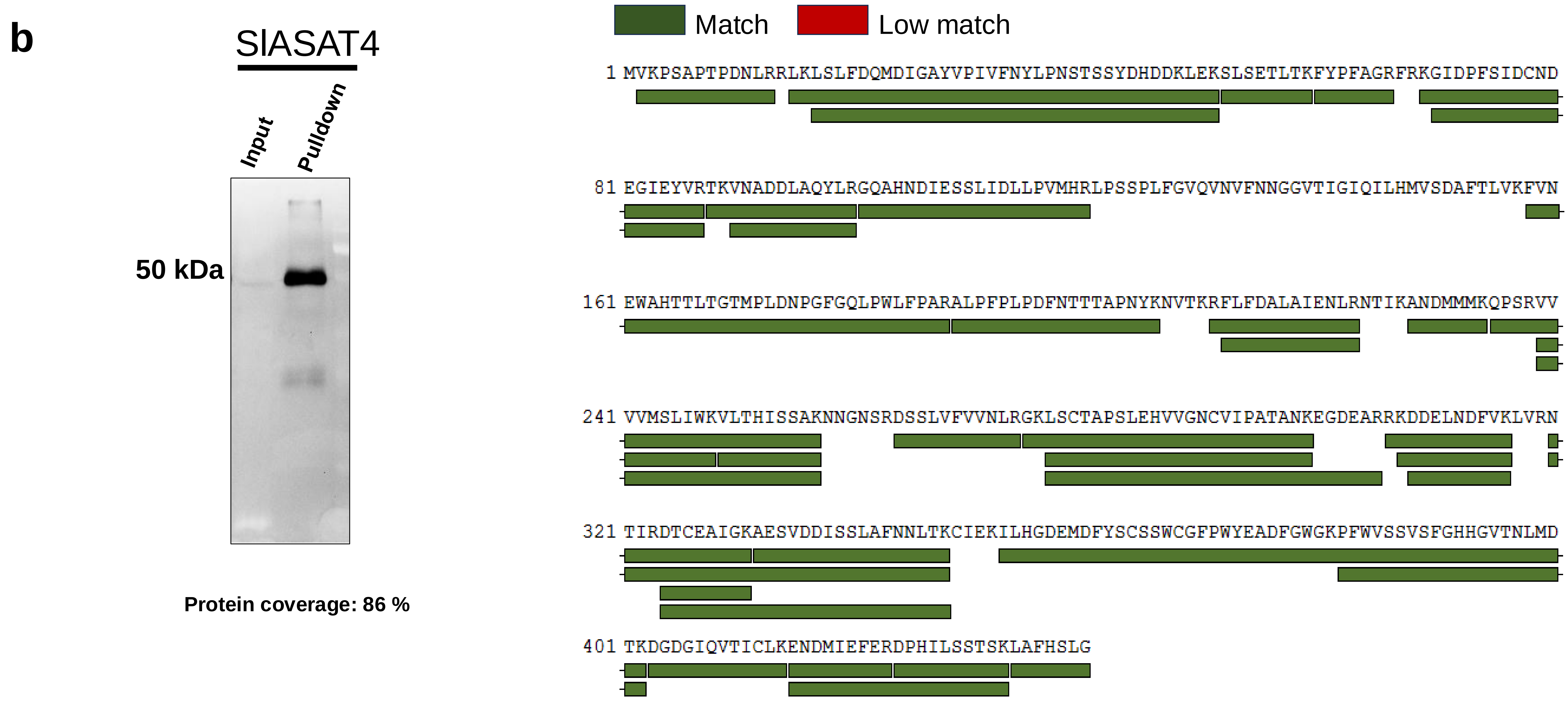


**Supplementary Fig. 9 SlASAT3 and SlASAT4 pulldown optimization.** Protein gel blot and proteomics analysis of SlASATs pulldown. The protein gel blot analysis of the input and HA-SlASAT3 (**a**) and Flag-SlASAT4 (**b**) immunoprecipitation (IP) samples, along with the schematic overview of the SlASAT proteins detected using LC/MS-MS analysis. IP was performed with anti-HA or Flag antibody. The experiment was repeated at least three times, with similar results. The peptides identified in the LC/MS analysis are represented by green boxes (high match > 90% amino acid identity), while the low match peptides are represented by red boxes (< 90% amino acid identity).

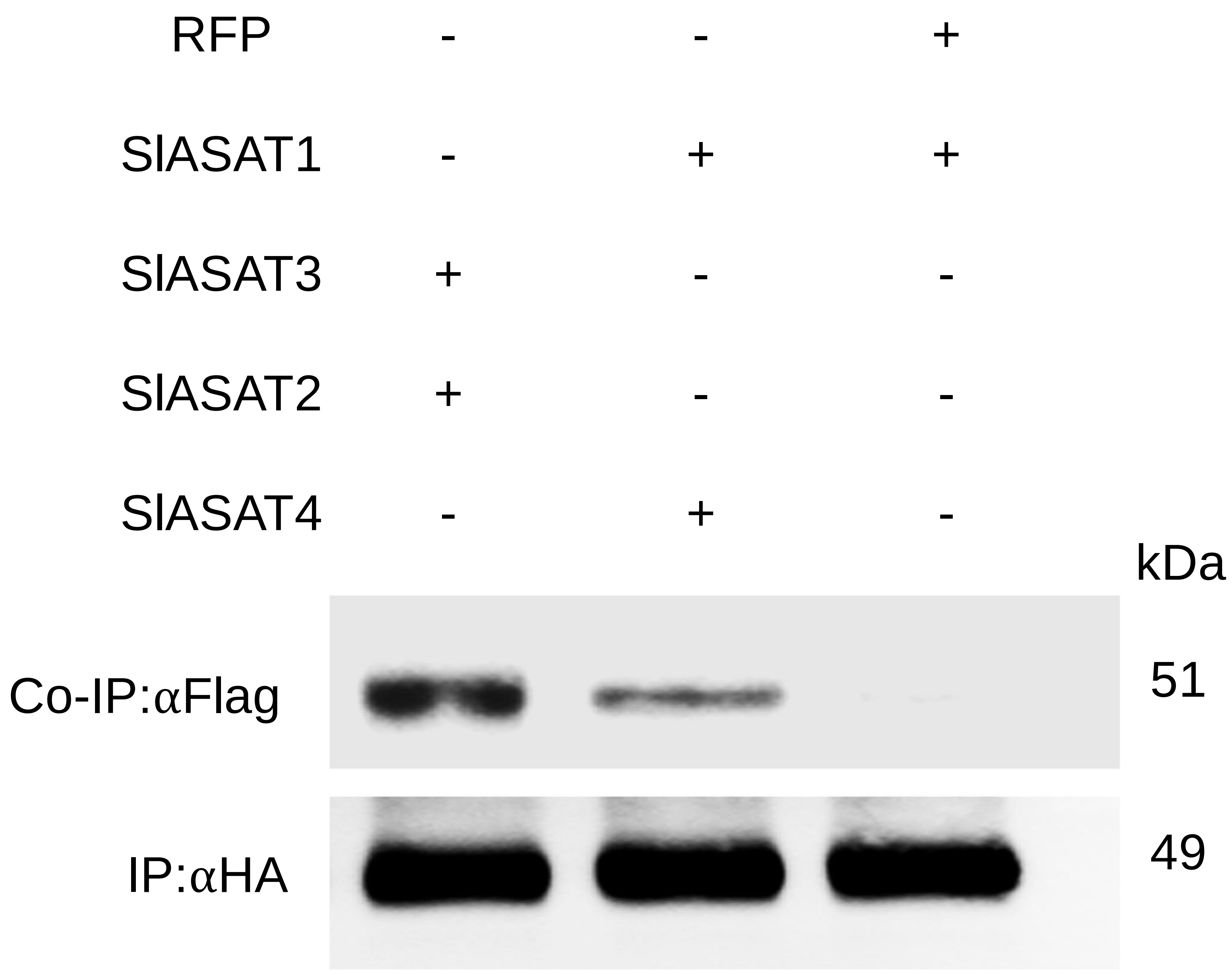


**Supplementary Fig. 10 Co-immunoprecipitation (Co-IP) showed specific interactions among SlASAT proteins.** HA- and Flag-tagged fusion proteins were transiently expressed in *N. benthamiana* leaves, and total proteins were extracted 3 days after infiltration. Protein extracts were prepared from *N. benthamiana* leaves transiently expressing combination of 35S::HA-SlASAT1, 35S::Flag-SlASAT2, 35S::HA-SlASAT3, 35S::Flag-SlASAT4 and 35S::Flag-RFP. Protein gel blot analysis samples immune precipitated (IP) with HA. IP was performed with anti-HA antibody and interacting proteins were analyzed with an anti-Flag antibody. The experiment was repeated at least three times, with similar results.

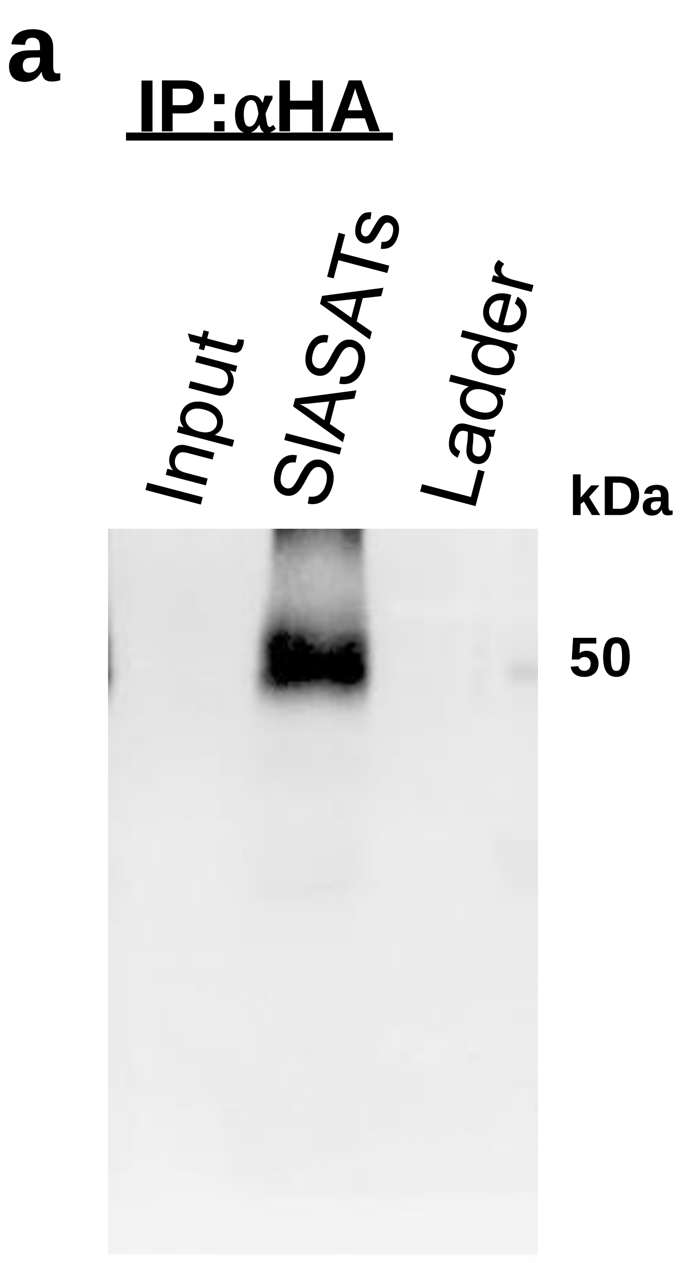


**Supplementary Fig. 11 Immunoprecipitation (IP) analysis of co-expressed SlASAT1-4 proteins.** HA-tagged SlASAT1 and Flag-tagged SlASAT4 were infiltrated in *N. benthamiana* together with untagged SlASAT2 and SlASAT3. Protein extracts were prepared from *N. benthamiana* leaves transiently expressing all four SlASATs. **a** Protein gel blot analysis of input and HA-IP samples. IP was performed with anti-HA magnetic beads. **b** Protein gel blot analysis of input and Flag-IP samples. IP was performed with anti-Flag magnetic beads. The experiment was repeated at least three times, with similar results.

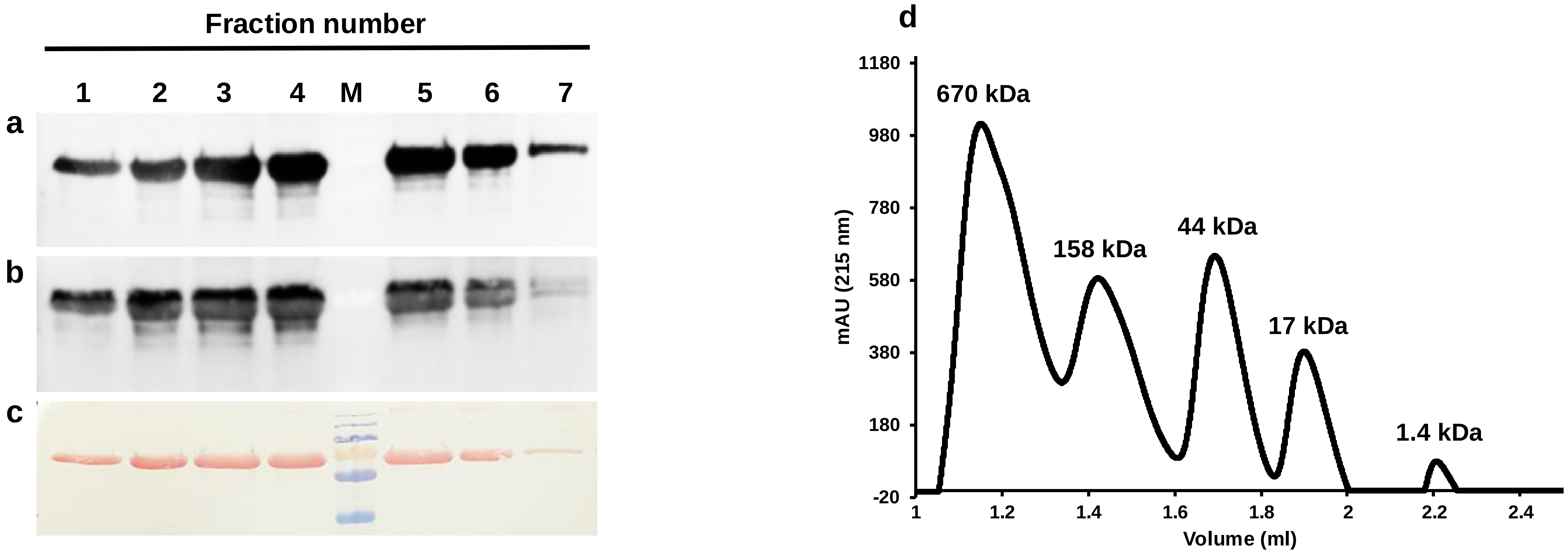


**Supplementary Fig. 12 Analysis of the size exclusion chromatography of the SlASAT pulldown complex.** Protein extracts were prepared from *N. benthamiana* leaves transiently expressing a combination of 35S::HA-SlASAT1, 35S::SlASAT2, 35S::SlASAT3, and 35S::Flag-SlASAT4, the pulldown complex was eluted using non-denaturing conditions with HA peptide and separated over size exclusion chromatography (SEC). **a** Protein gel blot analysis using anti-HA antibody of various SEC fractions showing the abundance of SlASAT1. **b** Protein gel blot analysis using anti-Flag antibody of various SEC fractions showing the abundance of SlASAT4. **c** The gel blot was stained with Ponceau S. **d** A set of globular protein standards was separated using the identical size exclusion column and method as used in Fig. 5a for the separation of the SlASAT1-4 complex. The molecular weights of the protein standards are indicated in kDa above each peak. The protein standards were monitored using a 215 nm wavelength.

| Group- ID | Protein Descriptions |
|---|---|
| NbACAT1a | acetyl-CoA C-acetyltransferase activity |
| E5LLE7 | Phosphoglycerate kinase |
| NbG6PDH | glucose-6-phosphate dehydrogenase activity |
| NbPDR2a | ABC-type transporter activity |
| NbPDR2b | ABC-type transporter activity |
| NbMVD1a | isoprenoid biosynthetic process |
| NbRanBP1-1a | nucleocytoplasmic transport |
| A0A060IKL | fatty acid biosynthetic process |
| A0A1U9IKY9 | mitochondria-associated ER membrane contact site |
| A0A288UJC0 | acyltransferase activity |

**Supplementary Fig. 13 Proteomics analysis of SlASAT1-4.** The List of top candidates identified in the proteomics analysis of the SlASAT1-4 metabolic complex. The top candidates were those that were common across all replicates and had the highest abundance
